# Exploiting Bacterial Effector Proteins to Uncover Evolutionarily Conserved Antiviral Host Machinery

**DOI:** 10.1101/2024.01.29.577891

**Authors:** Aaron Embry, Nina S. Baggett, David B. Heisler, Addison White, Maarten F. de Jong, Benjamin L. Kocsis, Diana R. Tomchick, Neal M. Alto, Don B. Gammon

## Abstract

Arboviruses are a diverse group of insect-transmitted pathogens that pose global public health challenges. Identifying evolutionarily conserved host factors that combat arbovirus replication in disparate eukaryotic hosts is important as they may tip the balance between productive and abortive viral replication, and thus determine virus host range. Here, we exploit naturally abortive arbovirus infections that we identified in lepidopteran cells and use bacterial effector proteins to uncover host factors restricting arbovirus replication. Bacterial effectors are proteins secreted by pathogenic bacteria into eukaryotic hosts cells that can inhibit antimicrobial defenses. Since bacteria and viruses can encounter common host defenses, we hypothesized that some bacterial effectors may inhibit host factors that restrict arbovirus replication in lepidopteran cells. Thus, we used bacterial effectors as molecular tools to identify host factors that restrict four distinct arboviruses in lepidopteran cells. By screening 210 effectors encoded by seven different bacterial pathogens, we identify six effectors that individually rescue the replication of all four arboviruses. We show that these effectors encode diverse enzymatic activities that are required to break arbovirus restriction. We further characterize *Shigella flexneri*-encoded IpaH4 as an E3 ubiquitin ligase that directly ubiquitinates two evolutionarily conserved proteins, SHOC2 and PSMC1, promoting their degradation in insect and human cells. We show that depletion of either SHOC2 or PSMC1 in insect or human cells promotes arbovirus replication, indicating that these are ancient virus restriction factors conserved across invertebrate and vertebrate hosts. Collectively, our study reveals a novel pathogen-guided approach to identify conserved antimicrobial machinery, new effector functions, and conserved roles for SHOC2 and PSMC1 in virus restriction.

**Author Summary:** Microbial pathogens such as viruses and bacteria encounter diverse host cell responses during infection. While viruses possess antagonists to counter these responses in natural host species, their replication can be restricted in unnatural host cells where their antagonists are ineffective. Bacteria also employ a diverse repertoire of immune evasion proteins known as “effectors” that can inhibit antimicrobial responses found in invertebrate and vertebrate hosts. In this study, we hypothesized that some bacterial effectors may target host immunity proteins that restrict both bacteria and viruses. To test this hypothesis, we screened a bacterial effector library comprising 210 effectors from seven distinct bacterial pathogens for their ability to rescue the replication of four viruses in insect cells that are normally non-permissive to these viruses. Though numerous effectors were identified to rescue the replication of each virus, the uncharacterized IpaH4 protein encoded by the human pathogen *Shigella flexneri* was able to rescue all four viruses screened. We discovered that IpaH4 enhances arbovirus replication in both restrictive insect and permissive human cells by directly targeting two novel, evolutionarily conserved antiviral host proteins, SHOC2 and PSMC1, for degradation. Our study establishes bacterial effectors as valuable tools for identifying critical antimicrobial machinery employed by eukaryotic hosts.

## Introduction

Arboviruses comprise a diverse group of <u>ar</u>thropod-<u>bo</u>rne viruses that are transmitted by dipteran (fly and mosquito) vectors to animal and human hosts. For example, vesicular stomatitis virus (VSV) is a negative-sense single-stranded (ss)RNA virus belonging to the *Rhabdoviridae* family that is the leading cause of vesicular disease in livestock in the United States, resulting in costly animal quarantines and trade embargoes [1]. Of the ∼500 arboviruses that have been identified, ∼150 are known to cause disease in humans [2]. Consequently, in 2022, the “Global Arbovirus Initiative” was launched by the World Health Organization to monitor and control arboviral disease [3]. Notable among arboviruses causing disease in humans are the positive- sense ssRNA viruses belonging to the *Togaviridae* family. This family includes chikungunya virus, the second-most prevalent arbovirus infecting humans worldwide [2]. However, the need for biosafety level-3 facilities to culture wild-type strains of chikungunya virus poses significant challenges to studying this togavirus. In contrast, other less pathogenic togaviruses [e.g. Ross River virus (RRV), O’nyong’nyong virus (ONNV), Sindbis virus (SINV)], can be cultured under BSL-2 conditions and thus have become important models for understanding togavirus-host interactions [4, 5]. However, we still lack vaccines and antiviral drugs to combat most human arbovirus infections, including those caused by togaviruses [5]. Thus, the identification of immune mechanisms that restrict arbovirus replication may provide additional avenues for the development of effective strategies to combat arboviral disease.

While genome-wide CRISPR-Cas9 and RNA interference (RNAi) screening platforms have been used to identify host immunity factors affecting arbovirus replication [6–8], these assays can be difficult, time-consuming, and cost prohibitive to set up, and are not easily applicable to non-model host systems. Moreover, these assays cannot provide insight into the strategies used by pathogens to combat host antiviral factors identified in these screens. Identification of pathogen-encoded immune evasion proteins (IEPs) targeting host immunity factors is important for several reasons. First, the existence of such IEPs is strong evidence for the physiologic importance of these interactions during the “molecular arms race” between pathogen and host. Second, while some IEPs simply bind/sequester host factors to inhibit their function, others can alter post-translation modifications to modify stability or function [9]. Thus, IEPs can be used as “tools” to both identify the host immunity factors they target and uncover molecular mechanisms that regulate host factor function. Third, IEPs often drive virulence and thus their characterization can reveal pathogenesis mechanisms [10, 11]. It is paramount to develop simplistic, functional assays that can both identify key antiviral factors restricting viral replication and that provide molecular tools to mechanistically dissect the function of such immunity factors.

Although arboviruses are well-adapted to replicate in dipteran and mammalian hosts, we have previously shown that several arboviruses, such as VSV and SINV, undergo abortive infections in cells derived from lepidopteran (moth and butterfly) hosts [12, 13]. For example, in *Lymantria dispar* (spongy moth)-derived LD652 cells, VSV and SINV undergo abortive infections post-entry after limited gene expression. However, their replication can be rescued by global inhibition of host transcription or by expression of mammalian poxvirus-encoded IEPs termed “A51R proteins”, suggesting that innate antiviral defenses block VSV and SINV replication in LD652 cells [12]. However, the host immune responses that are at play during restricted arboviral infections in LD652 cells remain poorly defined. More recently, we have reported the full genomic sequence of *L. dispar* and the LD652 cell transcriptome [14], making virus-LD652 cell systems more amenable to uncovering pathogen-host interactions at the molecular level. Our finding that mammalian poxviral IEPs can retain immunosuppressive function in LD652 cells suggests that some pathogen-encoded IEPs target host machinery conserved between insects and mammals. Thus, we were interested in identifying IEPs from other mammalian pathogens that promote arbovirus replication in LD652 cells. Such IEPs might be useful molecular tools in identifying the conserved host immunity factors they target.

Bacterial pathogens encode a wide array of IEPs that can manipulate eukaryotic immune responses. Many of these bacterial IEPs are “effector” proteins that are injected into eukaryotic host cells through bacterial secretion systems [10, 15]. These effectors can manipulate, usurp, and/or inhibit a variety of cellular processes once inside the host cell cytoplasm including cytoskeletal dynamics, host signaling cascades, and innate immune responses [10, 15]. Interestingly, some bacterial effectors inhibit innate immune pathways that are also antagonized by viruses, such as the Type I interferon (IFN) response [16], suggesting that bacterial and viral pathogens may need to evade common eukaryotic defense mechanisms. Although significant advances have been made towards understanding effector biology, the function of many effectors remains unknown. Understanding bacterial effector function is important because these proteins can be critical drivers of bacterial pathogenesis [10, 15]. However, defining the role of individual effectors during infection can be challenging due to functional redundancy among independent effectors encoded by a single bacterial pathogen [17]. Therefore, experimental strategies to study effector functions outside of bacterial infections may be useful for determining their role during natural infection.

Here, we further explore the restricted infections of arboviruses in lepidopterans by infecting moth cells with the rhabdovirus, VSV, and the togaviruses: SINV, RRV, and ONNV. We develop a simple, yet innovative approach to uncover evolutionarily conserved antiviral factors through the identification of bacterial effectors that rescue arbovirus replication in LD652 cells. By expressing a library of 210 effector proteins encoded by seven distinct bacterial pathogens, we identify six effectors capable of rescuing all four restricted arboviruses in LD652 cells: SopB, IpgD, HopT1-2, HopAM1, Ceg10, and IpaH4. Using mutagenesis, we demonstrate the importance of diverse enzymatic functions for SopB, IpgD, HopAM1, and IpaH4 in breaking arbovirus restriction. Moreover, crystallography and cell cultures studies reveal Ceg10 to encode a putative cysteine protease function that is required for arbovirus rescue. By focusing on the *Shigella flexneri*- encoded effector IpaH4, we reveal this novel bacterial E3 ubiquitin ligase to directly target two conserved host proteins, SHOC2 and PSMC1, for degradation in moth and human cells. To our knowledge, roles for these host factors in virus restriction had not been reported in any eukaryotic system. However, we show that depletion of intracellular SHOC2 or PSMC1 levels in moth or human cells promotes arbovirus replication, suggesting they have ancient roles in combating viral infection across diverse eukaryotic host species. Together, our findings demonstrate the utility of using naturally abortive arbovirus infections in lepidopteran cells for the interrogation of arbovirus- host interactions and establish it as a model for identifying conserved host immunity proteins targeted by pathogens.

## Results

### Inhibition of Host Transcription Rescues Restrictive Arbovirus Replication in LD652 Cells

Previously, we showed that the normally abortive infection of VSV and SINV in LD652 cells can be rescued by treatment of cultures with actinomycin D (ActD), an inhibitor of transcription [12]. ActD globally blocks transcription by host DNA-dependent RNA polymerases by intercalating into GC-rich regions of cellular DNA and thus does not impede viral RNA- dependent RNA polymerase-mediated transcription [18, 19]. The relief of arbovirus restriction by ActD treatment suggests that VSV and SINV undergo abortive infections in LD652 cells due to cellular antiviral responses that require active transcription [12]. To confirm these previous results and to determine if ActD treatment could relieve restriction of additional togaviruses related to SINV, such as RRV and ONNV, LD652 cells were infected with GFP reporter viruses (VSV-GFP [12], SINV-GFP [12], RRV-GFP [20], and ONNV-GFP [21]) in the absence or presence of ActD. Cells were then stained with CellTracker™ Orange Dye and imaged 72 h post-infection (hpi). Representative GFP fluorescence images and quantitative GFP signals (normalized to cell number with CellTracker™ signals) were used as a readout for viral replication and are shown in **Figure 1AB**. As expected, 0.05 μg/mL ActD treatment increased GFP signal in VSV-GFP and SINV-GFP infections by ∼10,000- and 100-fold, respectively (**Figure 1B**). Additionally, ActD treatment during VSV-GFP and SINV-GFP infection increased their viral titer by ∼1,000-fold for both viruses (**Figure 1C**). During ONNV-GFP and RRV-GFP infections, ActD increased GFP signal and viral titer by ∼100-fold and ∼10-fold, respectively (**Figure 1ABC)**. Importantly, we have previously shown that LD652 cells treated with this dose of ActD retain ∼90% viability [12], and thus enhanced virus replication is not due to a general decrease in cell viability. Together, these findings suggest that arbovirus infection of LD652 cells induces a restrictive immune response that requires active host transcription.

**Figure 1.**
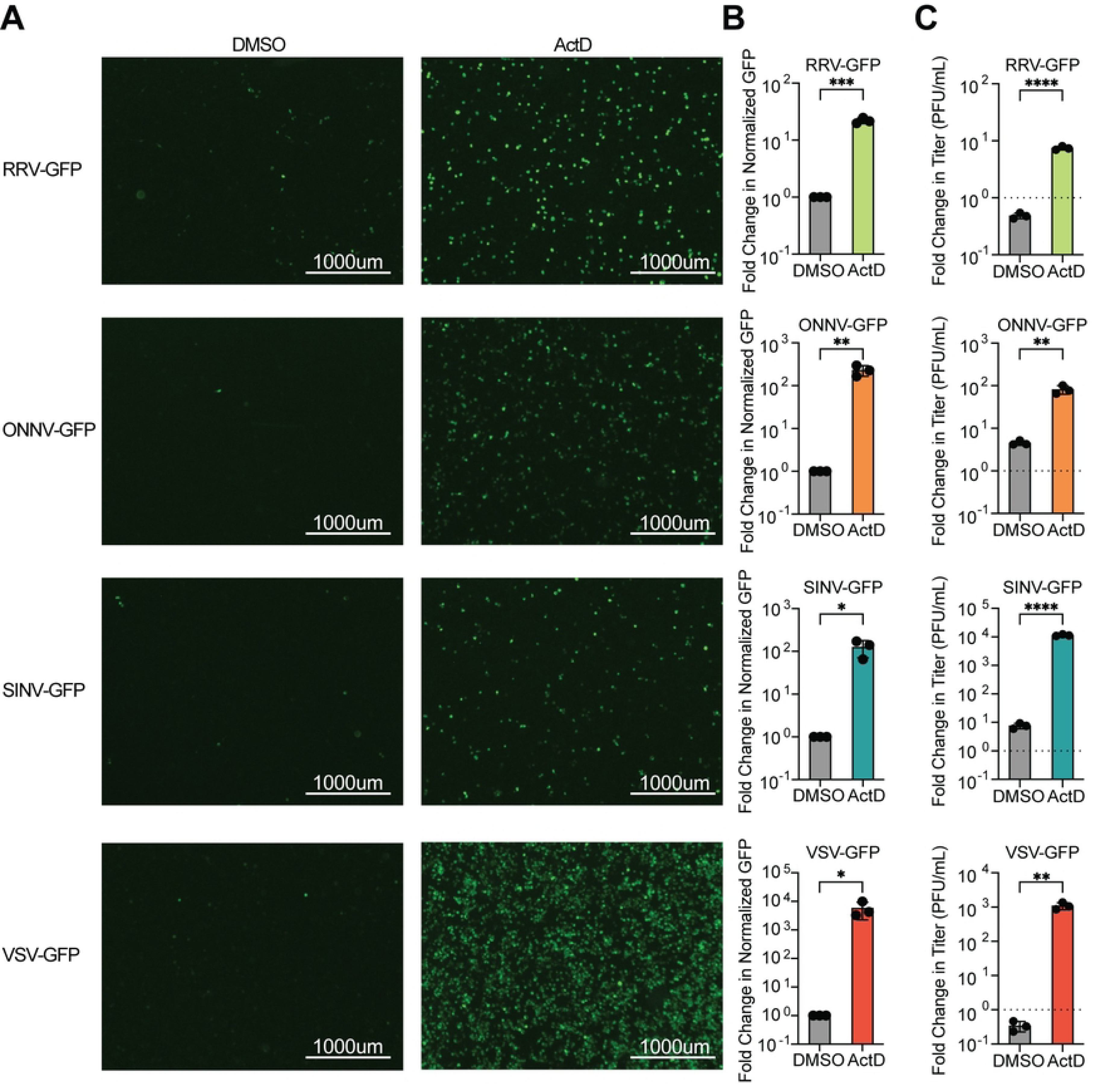
Abortive arbovirus replication in LD652 cells can be relieved with ActD treatment. **A.** Representative fluorescence microscopy images (GFP channel) of LD652 cells treated with DMSO (vehicle) or 0.05 µg/mL ActD and infected with the indicated GFP reporter strains for 72 h. **B.** Fold-change in normalized GFP signals in ActD-treated cultures relative to DMSO treatments. Cells were stained 72 hpi with CellTracker™ Orange dye (not shown) and imaged in GFP and RFP channels to calculate fold-change in GFP signal after normalization of cell number using CellTracker™ (RFP) channel signals. **C.** Fold-change in titer of supernatants from LD652 cell cultures treated as in **A-B** 72 hpi relative to input inoculum (dotted line). Data in B-C are means ± SD; n=3. Statistical significance was determined with unpaired student’s t-test; ns=P>0.1234, *=P<0.0332, **=P<0.0021, ***=P<0.0002, ****=P<0.0001.

### Specific Bacterial Effectors Relieve Arbovirus Restriction in LD652 Cells

We have previously shown that poxvirus-encoded A51R proteins are IEPs that rescue restricted arbovirus replication when expressed from plasmids transfected into LD652 cells [12, 22]. Therefore, we asked if bacterial effector proteins, which often function as IEPs, could also rescue arbovirus replication. To do this, we adapted and expanded a previously described effector library for expression in insect cells [23]. Briefly, 210 secreted bacterial effectors from seven pathogens (*Shigella flexneri*, *Salmonella enterica* serovar Typimurium, *Pseudomonas syringae*, Enterohemorrhagic *E. coli* O157:H7, *Yersinia pseudotuberculosis*, *Legionella pneumophila*, and *Bartonella henselae*) were cloned into the pIB/V5-His insect expression vector and screened for their ability to alleviate arbovirus restriction in LD562 cells. Cells were transiently transfected for 48 h and then infected with either GFP reporter viruses (RRV-GFP and ONNV-GFP) or luciferase reporter strains (SINV-LUC and VSV-LUC [12, 22]) for 72 h (**Figure 2A**). Effector proteins that enhanced viral GFP signals by >2.5-fold or luciferase signals by >4-fold above empty vector- transfected cells, were considered “hits” in our screen (**Figure 2B-E**). These cutoffs were chosen to avoid false positives stemming from experimental noise within our screening system and allowed us to focus on effectors that robustly rescued arbovirus replication. Of the 210 effector proteins screened, 10 effectors rescued RRV-GFP, 11 rescued ONNV-GFP, 18 rescued SINV- LUC, and 10 rescued VSV-LUC (**Figure 2B-F** and **Table S1**). Interestingly, effectors generally rescued in a virus-specific manner. For instance, 21 effectors only rescued one of the four arboviruses screened (**Figure 2F**). This suggests that these effectors may relieve virus-specific restrictions to replication. In contrast, seven effectors rescued three or more arboviruses: IpaH4, SopB, HopT1-2, HopAM1, Ceg10, EspK, and SidM (**Figure 2F**), suggesting that these effectors may target host restriction mechanisms that are active against a broader range of viral pathogens.

**Figure 2.**
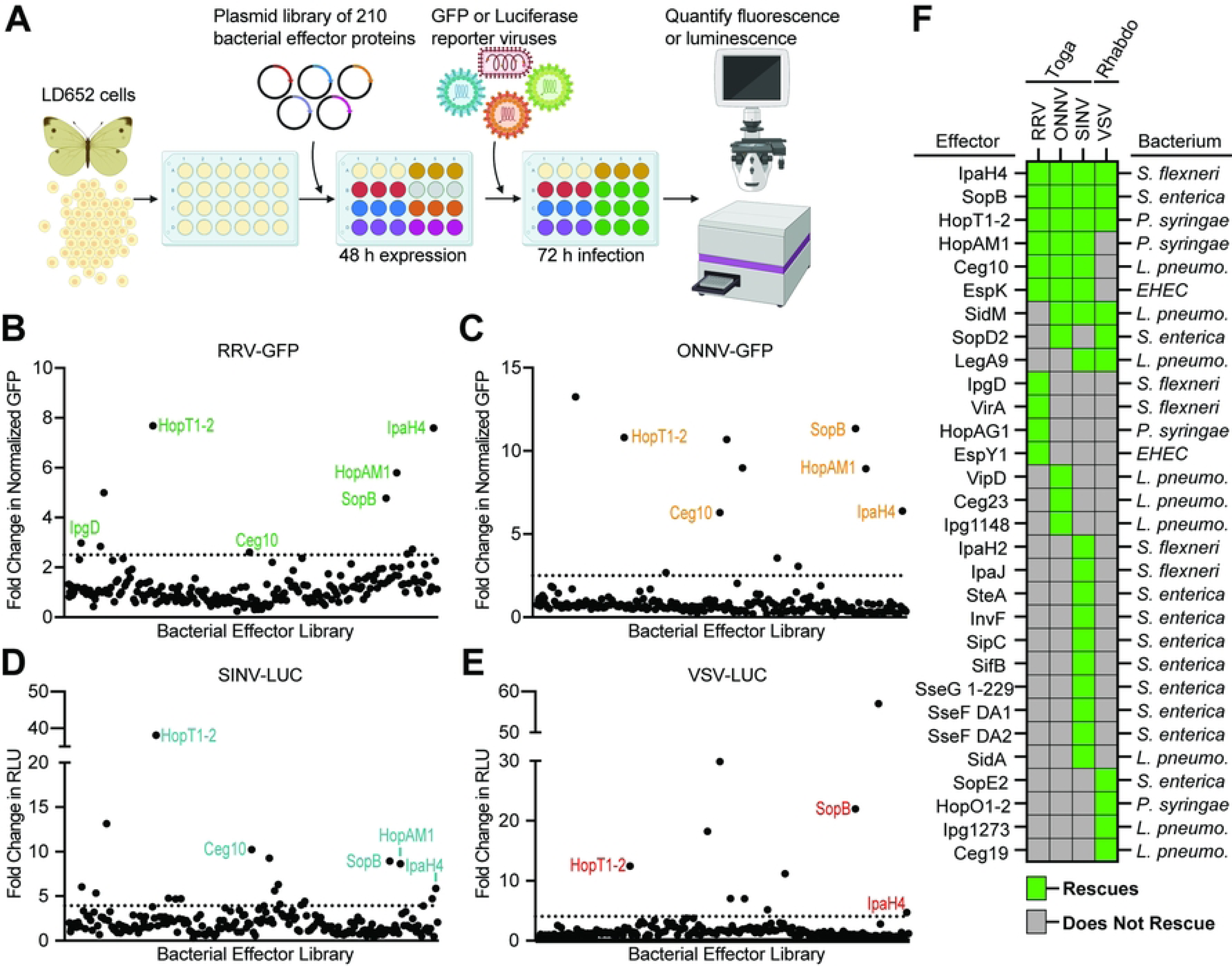
Specific Bacterial Effectors Relieve Arbovirus Restriction in LD652 Cells. A. Schematic outlining screen for bacterial effectors that rescue arbovirus restriction in LD652 cells. Cells were transfected with expression plasmids from a library consisting of 210 different effector proteins. After 48 h, cells were infected with either GFP or luciferase reporter strains. At 72 hpi, viral replication was quantified using fluorescence microscopy (RRV-GFP and ONNV-GFP) or luciferase assays (VSV-LUC and SINV-LUC). Image was created with Biorender.com. **B-E**. Fold- change in reporter readout, normalized to empty vector controls for all four screens. The cutoff for fold-change in GFP-based assays was set to >2.5, while the cutoff for luciferase reporters was set to >4-fold (represented by dotted horizontal lines). Data points are means. RLU = relative light units. **F.** Summary of bacterial effector proteins that rescued at least one virus. Green blocks indicate the effector rescued the virus indicated in the column header. The bacterium encoding each effector is noted to the right: *Shigella flexneri* (*S. flexneri*), *Pseudomonas syringae* (*P. syringae*), *Salmonella enterica* (*S. enterica*), *Legionella pneumophila* (*L. pneumo.*) *Enterohemorrhagic Escherichia coli* 0157:H7 (EHEC). Additional effector proteins from *Yersinia pseudotuberculosis* and *Bartonella henselae* were also screened but did not rescue arbovirus replication.

### Diverse Effector Activities are Required for Viral Rescue

Because our original effector library did not encode epitope-tagged effector genes, we wanted to confirm the expression and rescue functions of key hits from our screen. To do this, we chose five effectors: SopB, HopT1-2, HopAM1, Ceg10, and IpaH4 that rescued at least 3/4 arboviruses screened. These five effectors are collectively encoded by four bacterial pathogens and have distinct known or putative structures and functions (**Figure 3A**). We cloned these five effectors into a novel expression vector, pDGOpIE2, along with Flag epitope tags (**Figure 3B**). The pDGOpIE2 vector uses the same baculovirus-derived OpIE2 promoter [24] to drive effector expression as in the pIB/V5-His vector used in our initial screen, but the former vector encodes unique restriction sites that facilitated restriction enzyme-based cloning (**Figure S1**). Using these constructs, we confirmed the expression of all five Flag-tagged effectors by immunoblot in LD652 cells. We also confirmed expression of specific point mutants in some of these effectors (described below) predicted to inactivate effector enzymatic activities (**Figure 3B**). We next transfected either wild-type or mutant effector expression constructs into LD652 cells to determine: 1) if these Flag-tagged effectors could rescue arbovirus replication and 2) if known or putative enzymatic functions were required for viral rescue. Below we discuss each of these five effectors and their ability to rescue restricted arbovirus replication.

**Figure 3.**
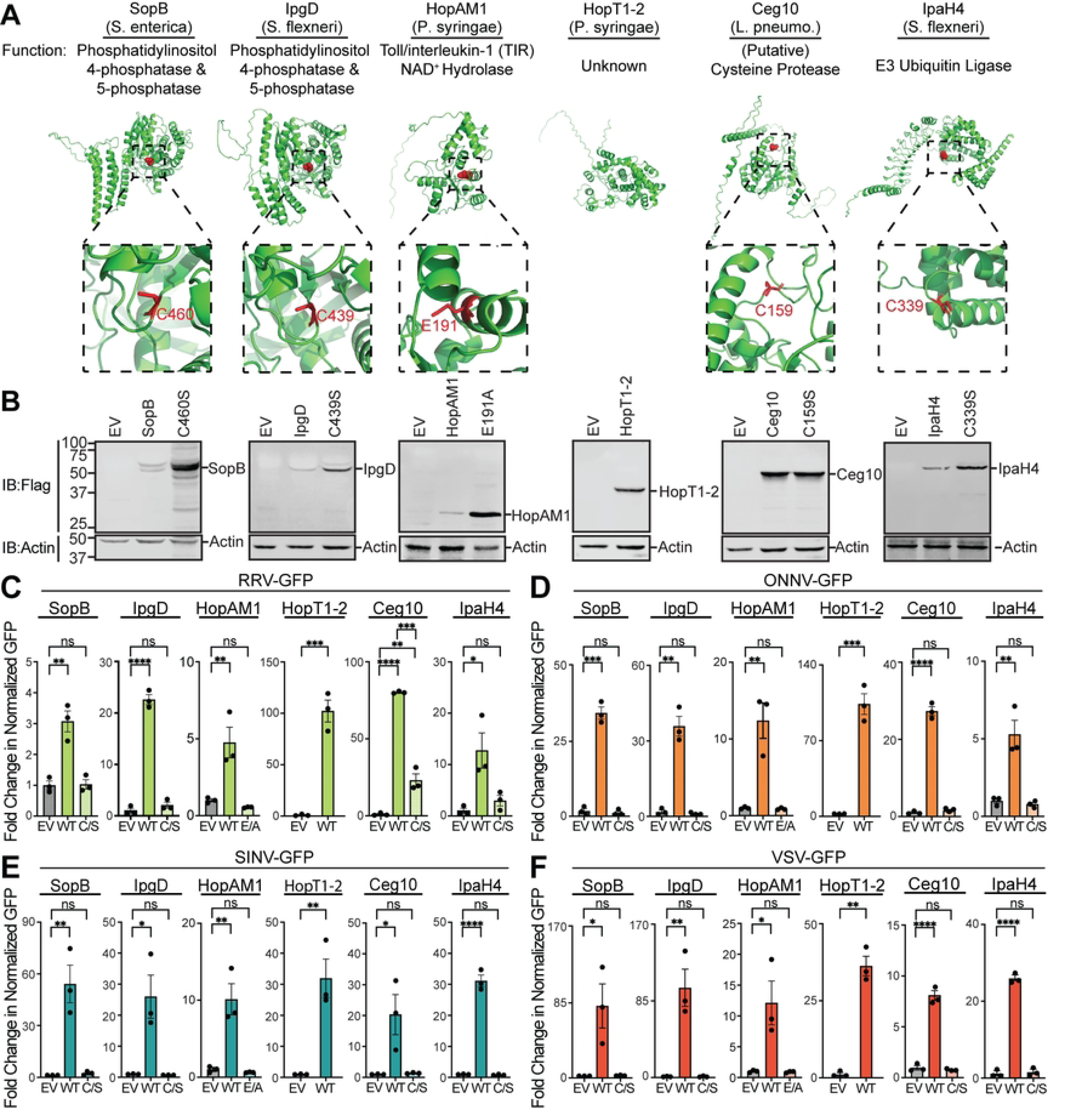
Validation and characterization of top hits from bacterial effector screens. A. AlphaFold predicted structures, and known or predicted enzymatic functions for indicated effector proteins identified as hits in arbovirus screens. Effector catalytic residues where substitution mutations were made are highlighted in red in AlphaFold structures. **B.** Representative immunoblots of Flag-tagged bacterial effector expression in LD652 cells 48 h post transfection. **C-F.** Fold-change in normalized viral GFP signal relative to empty vector (EV) controls 72 hpi with after transfection with indicated Flag-tagged effector constructs. Wild-type (WT) effectors are compared to their mutants (E/A or C/S). Cells were stained 72 hpi with CellTracker™ dye and imaged to calculate fold-change in normalized GFP signal over signals in EV treatments. Data in C-F are means ± SD; n=3. Statistical significance was determined with unpaired student’s t-test; ns=P>0.1234, *=P<0.0332, **=P<0.0021, ***=P<0.0002, ****=P<0.0001.

SopB is encoded by *Salmonella enterica* serovar Typimurium and functions as a phosphatase to generate phosphatidylinositol-5-phosphate (PI(5)P) from PI(4,5)P to activate Phosphoinositide 3-kinase (PI3K) signaling [25]. This activity has been shown to promote bacterial entry by inducing membrane ruffling [26] as well as altering endosomal maturation and trafficking [27]. The phosphatase activity of SopB relies on a catalytic cysteine (C420) [25] and mutants encoding a C420S substitution (SopB^C420S^) have impaired enzymatic activity [28, 29]. When expressed in LD652 cells, we found wild-type SopB to significantly enhance the replication of all four arboviruses, while SopB^C420S^ did not significantly affect arbovirus replication when compared to empty vector control treatments (**Figure 3C-F**). Importantly, this lack of rescue by the SopB^C420S^ mutant was not due to poor expression as it expressed to higher levels than wild- type SopB (**Figure 3B**). Interestingly, the degree of arbovirus rescue was variable among the viruses assayed. For instance, while SopB enhanced RRV-GFP by only ∼three-fold over empty vector controls, it increased the other three viruses by ∼30-80-fold. Although the absolute fold changes in arbovirus replication varied between our initial screen using pIB/V5-His and our confirmatory screen with pDGOpIE2, the overall trends in SopB rescue were similar (**Figure 2B- E** versus **Figure 3C-F**). Interestingly, the *Shigella flexneri*-encoded IpgD effector shares ∼47% amino acid identity to SopB and can also generate (PI(5)P) from PI(4,5)P reliant on a conserved catalytic cysteine (C439) [30, 31]. Therefore, it was surprising that IpgD only rescued one arbovirus in our initial screens. However, several immunoblot experiments failed to detect Flag- tagged IpgD expression in LD652 cells when cloned into pDGOpIE2, suggesting that IpgD may express more poorly in insect cells than SopB. Thus, we cloned a codon-optimized, Flag-tagged form of IpgD into pDGOpIE2 vectors and re-tested its ability to rescue all four arboviruses. Strikingly, we found the codon-optimized IpgD expressed in immunoblots (**Figure 3B**) and enhanced the replication of all four arboviruses (**Figure 3C-F**). This illustrates that poor expression of some effectors in our library may have produced false negatives. Importantly, expression of a catalytically-inactive mutant IpgD^C439S^ [32] failed to rescue arbovirus replication, despite robust expression (**Figure 3B-F**). These data suggest that a common phosphatase activity encoded by effectors from *S. enterica* and *S. flexneri* can break arbovirus restriction in LD652 cells.

The effector gene library used to screen for viral rescue included effectors encoded by the plant pathogen *Pseudomonas syringae*. It was therefore notable that two effectors, HopAM1 and HopT1-2, rescued arbovirus replication within the insect cells. These data suggest that host substrates of effectors, highly conserved through to Plantae, can reveal novel points of viral restriction. The HopAM1 effector possess a toll-like receptors and interleukin-1 receptors (TIR) domain that suppresses pattern-triggered immunity in plant hosts [33]. HopAM1 was recently shown to catalyze the formation of a novel cyclic adenosine monophosphate (ADP)-ribose (cADPR) isomer, termed “v2-cADPR” [33], which is required for its immunosuppressive function and for *P. syringae* pathogenicity [34]. The hydrolase activity of HopAM1 requires a catalytic glutamate (E191) to hydrolyze nicotinamide adenine dinucleotide (NAD^+^) into v2-cADPR [33]. Expression of Flag-tagged HopAM1 constructs in LD652 cells significantly enhanced the replication of all four arboviruses, while a catalytically-inactive mutant (HopAM1^E191A^) [35] did not significantly affect arbovirus replication (**Figure 3B-F**). HopT1-2 has also been implicated in suppressing plant immunity. Previous work has demonstrated that HopT1-2 antagonizes “nonhost resistance” defenses in plants [36] by suppressing expression of nonhost 1 proteins required for this immune response [37]. However, the molecular mechanism underlying this HopT1-2 function remains unknown. AlphaFold predicted structures (**Figure 3A**) and HHpred homology determination software [38] were unable to identify a putative catalytic activity for HopT1-2. Expression of Flag-tagged HopT1-2 was confirmed by immunoblot in LD652 cells and this construct rescued all four restricted arboviruses as we observed in our original screen (**Figure 3B-F**). Together, these results illustrate that two specific effectors encoded by a plant pathogen can antagonize antiviral responses in animal cells. Future studies will be needed to determine how HopAM1 generation of v2-cADPR breaks viral restriction, and the functional role of HopT1- 2.

In addition to identifying effector proteins with known host substrates, our unbiased screen uncovered novel effector proteins involved in breaking host immunity. Ceg10 is an uncharacterized effector secreted by *Legionella pneumophila*, the causative agent of human Legionnaires’ disease [39]. BLAST homology searches identified proteins closely related to Ceg10 in different *Legionella* serovars, yet sequence comparisons did not reveal a putative function. We therefore took a structural biology approach to help assign a function to Ceg10. We were unable to obtain crystals suitable for X-ray crystallography with full-length Ceg10, possibly owing to flexible N- and C-terminal domains as determined by the AlphaFold prediction (**Figure 3A**). We then used limited proteolysis to produce a core Ceg10 central domain (T55-R287) that was amenable to structural determination by crystallography (**Figure 4A**). We determined the structure of the trypsin limited-proteolysis protected core (Ceg10^TR^) from three different data sets to 1.7 Å, 1.4 Å and 1.5 Å resolution, respectively (**Table S2**).

**Figure 4.**
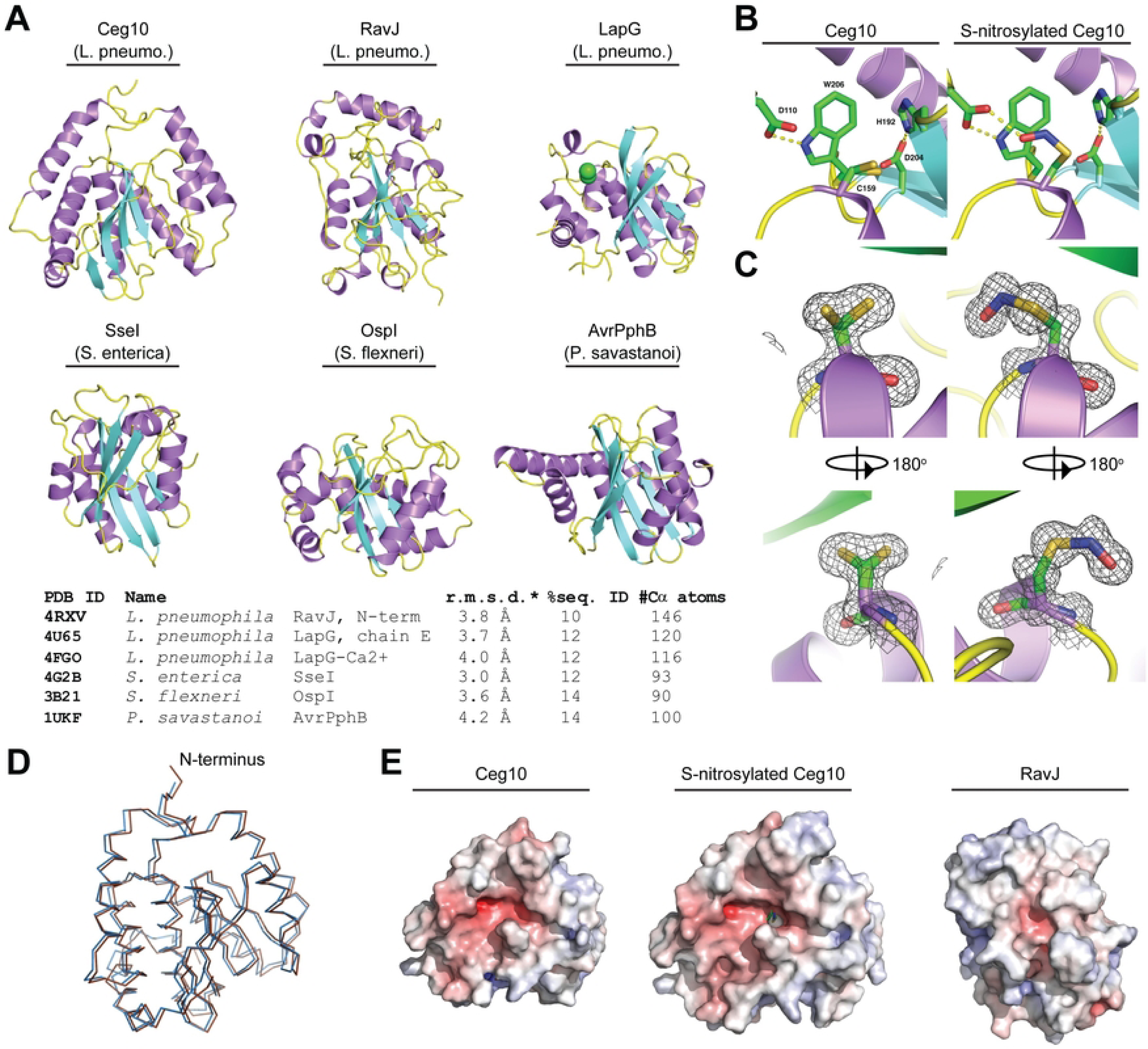
Structural analysis of *Legionella pneumophila* effector Ceg10. A. Six structural homologs of Ceg10, shown in the same orientation with the putative active site near the top of the figure. Top row, from left: Ceg10 (this study), *L. pneumophila* RavJ, *L. pneumophila* LapG. Bottom row, from left: *S. enterica* SseI, *S. flexneri* OspI, *P. savastanoi* AvrPphB. Table shows these structural homologs as determined via the Dali Lite server and their PBD ID**. B.** The putative active site of Ceg10 with residues shown in stick representation and hydrogen bonds are shown as dotted yellow lines. The catalytic Cys (C159), Asp (D204) and His (H192) are labeled as well as residues Asp110 and Trp 206 which are hydrogen bonded to each other in both structures. In the S-nitrosylated structure, Asp110 also hydrogen bonds to the nitrosylated-C159 and van der Waals interactions occur between the aromatic ring of Trp206 and the nitrosylation moiety. **C.** Electron density for C159 in the native (left) and S-nitrosylated (right) Ceg10 structures. The final refined 2Fo-mDFc electron density map, contoured at the 1σ level, is shown superimposed on each residue, as well as a 180° rotation of this region. **C.** Superposition of native (blue) and S- nitrosylated (brown) Ceg10 structures. **D.** Electrostatic surface potential of both Ceg10 structures and RavJ. All structures are orientated with the putative catalytic cysteine residue in approximately the center of the surface, and the orientations between Ceg10 and RavJ correspond to protein alignments. The displayed surface is colored by electrostatic potential from -10 kT (red) to + 10 kT (blue), as calculated by the APBS plugin in PyMOL.

Overall, Ceg10^TR^ is composed of 8 α-helices surrounding 7 β-sheets in a conformation resembling a cysteine proteinase with a putative catalytic triad consisting of Cys-159, His-196, and Asp-204 (**Figure 4AB**). A structural search via the PDBeFold [40] reveals several homologs containing the Cys-His-Asp triad but with low sequence identity (**Figure 4A**). Interestingly, while the Cys-159 (C159) of Ceg10 is modeled in two roughly equal conformers in data set 1, in data sets 2 and 3, C159 is modeled in only one conformer as a nitrosylated cysteine, or S-nitrosothiol [41] (**Figure 4BC**). Despite this, both structural determinations overlap modestly (**Figure 4D**). All crystals used for data sets in this study were grown from the same batch of purified protein, but the crystallization condition for data set 1 was 2.0 M sodium, potassium phosphate whereas for data sets 2 and 3 the major precipitants were polyethylene glycols. The significant level of trace divalent metal ions present in commercial preparations of sodium or potassium phosphate, and the extended time between initiation of crystal growth and harvesting is likely the reason the protein crystallized in data set 1 has lost the S-nitrosothiol, as metal catalyzed decomposition has been well documented for this post-translational modification [42]. In the S-nitrosothiol structure, the carboxylate of Asp-110 is hydrogen bonded to both the nitrosyl oxygen of C159 and the sidechain of Trp-206 (**Figure 4B**).

The closest homolog of Ceg10, the *Legionella pneumophila* effector RavJ, is the only homolog that includes an analogous tryptophan (Trp-172) and aspartate (Asp-24), but the cysteine (Cys-101) is neither modified nor modeled in multiple conformations. Analysis of the electrostatic potential mapped to the surface of Ceg10 and RavJ reveals that the catalytic site of RavJ is buried while the S-nitrosothiol of Cys-159 in Ceg10 is solvent exposed (**Figure 4E**). These data together indicate that Ceg10 harbors a highly reactive catalytic cysteine that may be critical for regulating host responses to infection. Indeed, expression of Flag-tagged wild-type Ceg10 significantly enhanced the replication of all four arboviruses. Consistent with our structural analysis, conversion of the predicted catalytic cysteine to serine (C159S) generated a mutant (Ceg10^C159S^) that was unable to rescue three of the four arboviruses screened, despite robust expression (**Figure 3B-F**). Although Ceg10^C159S^ significantly enhanced RRV-GFP replication compared to empty vector control, this rescue was significantly weaker than rescue observed with wild-type Ceg10 constructs (**Figure 3B-F**). Further studies will be needed to confirm Ceg10 function and its regulation by S-nitrosylation, however, these data suggest that a putative cysteine protease encoded by *L. pneumophila* is capable of targeting antiviral immune pathways in insect cells.

Lastly, IpaH4 is secreted by *Shigella flexneri* which causes Shigellosis in humans [43]. IpaH proteins are a family of bacterial E3 ubiquitin ligases that consist of two domains: a leucine rich repeat domain (LRR) used for substrate specificity and a novel E3 ligase domain (NEL) containing the catalytic cysteine [15, 44, 45]. We confirmed expression of wild-type IpaH4 and demonstrated significant viral rescue in all cases, however, an IpaH4 mutant with a serine substitution at position 339 (IpaH4^C339S^) was unable to rescue any arbovirus tested (**Figure 3B- F**). Our group [46–48] and others [49, 50] have shown that IpaH proteins antagonize various eukaryotic immune response pathways by targeting host defense proteins for degradation. However, IpaH4 has remained an uncharacterized member of the IpaH family. Thus, we were interested in both determining if IpaH4 was indeed an active E3 ubiquitin ligase and uncovering the putative host substrates that it targets.

### IpaH4 is an E3 ubiquitin Ligase that directly targets host PSMC1 and SHOC2 proteins

To confirm that IpaH4 exhibits E3 ubiquitin ligase activity, we performed *in vitro* autoubiquitination assays. E3 ubiquitin ligase activity can be revealed by demonstrating: 1) autoubiquitination function; 2) polymerization of free ubiquitin chains; and/or 3) direct ubiquitination of a substrate [51, 52]. Given that the substrates of IpaH4 are unknown, we conducted *in vitro* autoubiquitination experiments with GST-tagged IpaH4 (GST-IpaH4) purified from *E. coli*. Ubiquitination is a post-translation modification that transfers ubiquitin molecules from E1 (ubiquitin-activating enzymes) to E2 (ubiquitin-conjugating enzymes) to E3 ubiquitin ligases, which ultimately covalently attach ubiquitin to target proteins [53]. When mixed with the necessary components of a ubiquitination reaction: E1, E2, ubiquitin, and ATP, GST-IpaH4 displayed clear autoubiquitination as evidenced by the formation of higher molecular weight species that reacted with anti-GST and anti-ubiquitin antibodies (**Figure 5A**). However, when E1 was removed from these *in vitro* reactions, these higher molecular weight species were not detected, as expected (**Figure 5A**). Interestingly, under these reaction conditions, we were unable to observe autoubiquitination activity for two additional IpaH family members, IpaH2.5 and IpaH9.8 (**Figure 5A**). This finding is consistent with prior observations that IpaH members typically display autoinhibition unless in the presence of substrates [54, 55]. Thus, the robust IpaH4 autoubiquitination observed in the absence of substrates suggests an important distinction of IpaH4 autoregulation when compared with other IpaH members. GST-IpaH4 autoubiquitination activity could be detected at wide range of protein concentrations (0.2-1 μM) and typically became more obvious at higher IpaH concentrations. However, the GST-IpaH4^C339S^ mutant predicted to inactivate the catalytic cysteine, was unable to display detectable autoubiquitination even at 1 μM concentrations (**Figure 5B**). The robust autoubiquitination of wild-type IpaH4 may make it more prone to proteasomal degradation in host cells, and thus may contribute to the reduced expression we typically observe when we transfect wild-type and C339S mutant IpaH4 vectors into LD652 cells (**Figure 3B**).

**Figure 5.**
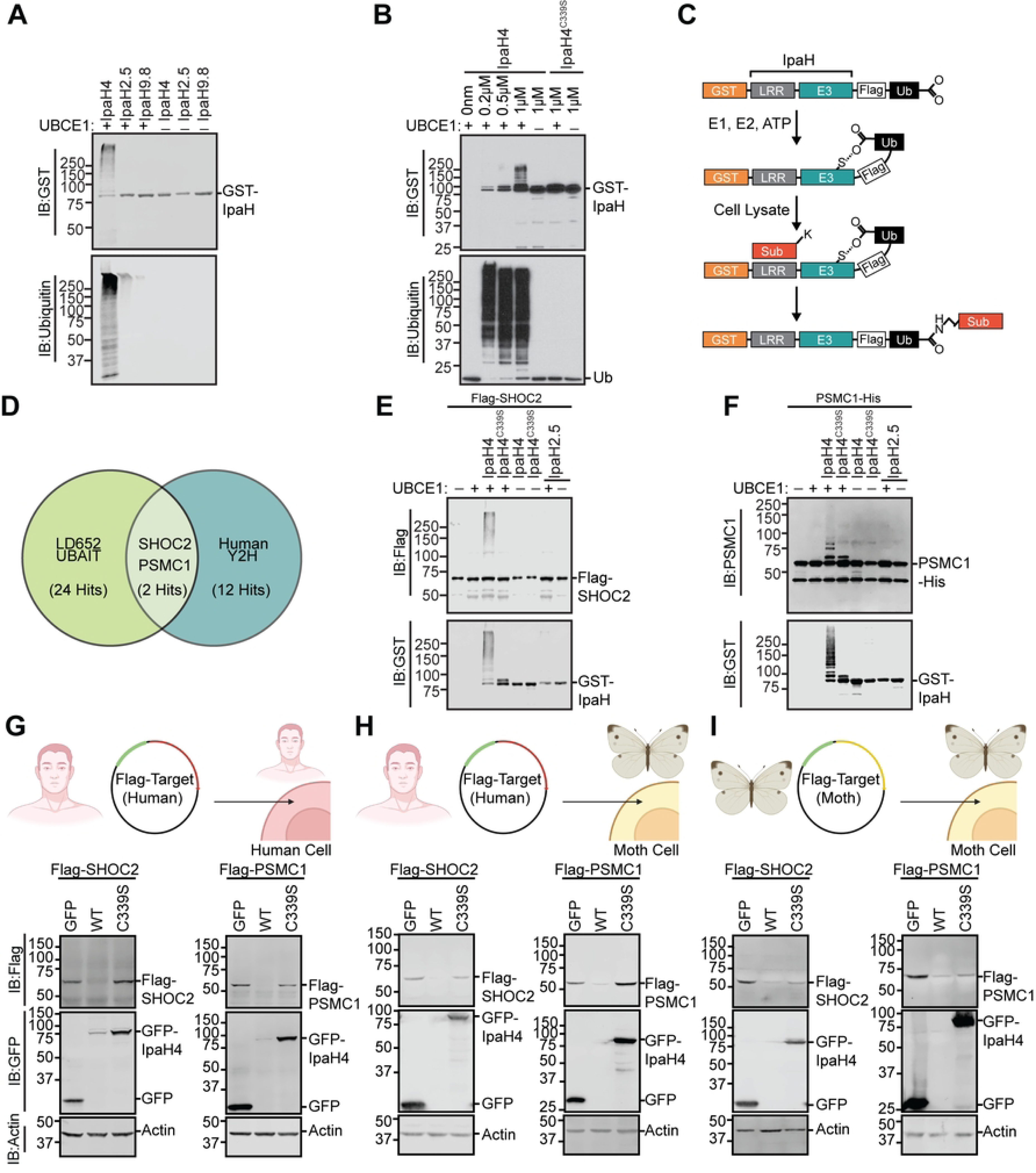
Identification of host SHOC2 and PSMC1 as conserved targets of IpaH4. A. Representative immunoblot of *in vitro* ubiquitination assay performed with indicated GST-IpaH proteins in the absence of substrates. **B.** Representative immunoblot of *in vitro* ubiquitination assay performed with indicated concentrations of wild-type GST-IpaH4 or GST-IpaH4^C339S^ mutant proteins. **C.** Schematic outlining UBAIT protocol [53, 56]. **D.** Venn diagram showing conserved putative substrates (overlapping region) of IpaH4 across UBAIT experiments (n=3) in LD652 cell lysates and Y2H screens (n=2) against a human prey library. **E.** Representative immunoblot of *in vitro* ubiquitination assay showing IpaH4-mediated ubiquitination of human Flag-SHOC2 proteins. **F.** Representative immunoblot of *in vitro* ubiquitination assay showing IpaH4-mediated ubiquitination of human PSMC1-His proteins. **G.** Representative immunoblot of degradation assays using indicated Flag-tagged human proteins in transfected HEK293T cells co-expressing GFP, IpaH4 (WT) or catalytic mutant GFP-IpaH4^C339S^ (C339S). **H.** Representative immunoblot of degradation assays using indicated Flag-tagged human proteins in transfected LD652 cells co- expressing GFP, IpaH4 (WT) or catalytic mutant GFP-IpaH4^C339S^ (C339S). **I.** Representative immunoblot of degradation assays of Flag-tagged moth (*L. dispar*) protein in LD652 cells expressing GFP, IpaH4 (WT) or catalytic mutant GFP-IpaH4^C339S^ (C339S). Image was created with Biorender.com.

Next, we sought to identify the host substrates of IpaH4. As IpaH4 is encoded by a human pathogen and was able to rescue arbovirus replication in insect cells, we hypothesized that the natural substrates of IpaH4 may be highly conserved between mammals and invertebrates. Therefore, we employed two distinct approaches to identify putative targets of IpaH4 in both moth and human backgrounds. First, ubiquitin activated interaction trap (UBAIT) assays [53, 56] were conducted in LD652 whole cell extract (**Figure 5C**). We have used this approach previously to identify the host substrates of other IpaH members [46, 47]. Briefly, the human ubiquitin gene was cloned in frame with IpaH4 to generate a C-terminal fusion protein (IpaH4^UBAIT^). This fusion allows IpaH proteins to bind their substrates through their N-terminal LRR domains and catalyze thiol-mediated ligation of the fused ubiquitin to a lysine on their substrates [56]. This results in the covalent linkage of IpaH proteins to their substrates, allowing their identification by mass spectrometry after affinity purification of IpaH-substrate complexes [56]. Using this approach, we identified 24 moth proteins that were enriched at least five-fold in IpaH4^UBAIT^ reactions compared to UBAIT reactions conducted with IpaH2.5^UBAIT^ constructs, which served as our control (**Figure 5D and Table S3**).

Because our goal was to identify evolutionarily conserved mechanisms of host immunity, we took advantage of an existing commercial human prey library and conducted two independent yeast two-hybrid (Y2H) screens using IpaH4 bait. An advantage of using a Y2H approach is that we can identify direct IpaH4-substrate interactions whereas UBAIT techniques tend to identify both direct and indirect IpaH-host interactions. Furthermore, by comparing the putative host substrates identified with UBAIT in the moth system with putative targets identified in human Y2H prey screens, we can detect substrate interactions common between two distinct approaches, and thus more likely to be bona fide interactors. Our Y2H screens with IpaH4 bait identified 12 [57] human proteins as putative IpaH4 interactors with >2 independent clones identified across two screens (**Figure 5D, Figure S2A and Table S3**). We then compared both UBAIT and Y2H screening results and found Leucine-rich repeat protein SHOC2 and proteasome 26S subunit, ATPase 1 (PSMC1; also known as Rpt2) to be the only two reproducibly identified putative substrates across moth and human backgrounds (**Figure S2A**).

To determine if IpaH4 directly ubiquitinates purified human SHOC2 or PSMC1 proteins, *in vitro* ubiquitination assays were conducted. Recombinant Flag-SHOC2 or PSMC1-His proteins were incubated with GST-IpaH4, E1, E2, ubiquitin, and ATP as described above for our autoubiquitination assays. Wild-type GST-IpaH4, but not GST-IpaH4^C339S^, was capable of ubiquitinating both host proteins in an E1-dependent manner, as evidenced by the formation of higher molecular weight Flag-SHOC2 and PSMC1-His species (**Figure 5E-F**). In contrast, incubation with GST-IpaH2.5 did not result in ubiquitination of either protein (**Figure 5E-F**). These results suggest that IpaH4 specifically ubiquitinates both SHOC2 and PSMC1 *in vitro*.

Lastly, given that other IpaH family members ubiquitinate substrates for proteasomal degradation [47, 49, 50], we hypothesized that intracellular SHOC2 and PSMC1 protein levels may be reduced in the presence of IpaH4. To assess this, degradation assays were conducted by co-expressing GFP-tagged IpaH4 with Flag-tagged human or moth SHOC2 and PSMC1 constructs in order to determine if IpaH4 activity altered intracellular levels of these putative substrates. Both human Flag-SHOC2 and Flag-PSMC1 levels were dramatically reduced when co-transfected with wild-type GFP-IpaH4 constructs, when compared to co-transfections with GFP- or GFP-IpaH4^C339S^-expressing vectors in HEK293T cells (**Figure 5G**). Other putative targets of IpaH4 (**Figure S2A**) such as SLU7, PRPF6, CRTC1, and BRD7 that were identified in our Y2H screen with human prey but not in our moth UBAIT assays, mostly showed little-to-no reduction in their levels in HEK293T cells when in the presence of GFP-IpaH4 (**Figure S2B**). One exception to this trend was human RNF214, which was dramatically reduced in level when co-expressed with GFP-IpaH4 (**Figure S2B**). However, RNF214 lacks a discernable *L. dispar* ortholog [14] and therefore was unlikely to be involved in arbovirus restriction in LD652 cells.

We next sought to determine if SHOC1 and PSMC1 were targeted by IpaH4 in the insect cell background. We found that human Flag-SHOC2 and Flag-PSMC1 proteins were dramatically reduced in abundance when co-transfected with GFP-IpaH4 in LD652 cells, suggesting that IpaH4 can target these human proteins when expressed in either human or moth cells (**Figure 5H**). To determine if IpaH4 can also alter the levels of moth-encoded SHOC2 and PSCM1 proteins, we co-transfected Flag-tagged versions of these putative moth targets into LD652 cells in the absence or presence of GFP-IpaH4. These experiments demonstrated that GFP-IpaH4, but not GFP-IpaH4^C339S^ constructs, could dramatically reduce moth Flag-SHOC2 and Flag- PSCM1 levels (**Figure 5I**). Collectively, these results suggest that wild-type, but not catalytically- inactive IpaH4 proteins, can reduce the levels of SHOC2 and PSMC1 proteins encoded by both *L. dispar* and human hosts.

### Depletion of SHOC2 and PSMC1 Rescues Restrictive Arbovirus Replication in LD652 Cells

One prediction of our approach is that the host substrates of effector proteins secreted by mammalian and plant pathogens are key regulators of viral restriction in the moth. To then determine if endogenous SHOC2 proteins contributed to arbovirus restriction in moth cells as our data suggests, we adapted a previously described CRISPR-Cas9 system for disrupting gene expression in lepidopteran insects [58] to inhibit SHOC2 expression. We cloned two independent single-guide RNAs (sgRNAs) targeting *L. dispar SHOC2* into pIE1-Cas9-SfU6-sgRNA-Puro [58], an “all-in-one” vector system that expresses Cas9 nuclease, sgRNA, and a puromycin resistance cassette for selection. As controls, cells were transfected with either empty vector or a sgRNA targeting the *L. dispar relish* gene, which encodes a Nuclear Factor-κB-like transcription factor that we have shown to contribute to VSV and SINV restriction in LD652 cells [12, 22]. Transfected cells underwent three rounds of puromycin selection and were then challenged with reporter arboviruses. As expected, cells expressing sgRNA targeting *relish* were significantly more susceptible to VSV-GFP and SINV-GFP infection when compared to empty vector control treatments. RRV-GFP and ONNV-GFP replication was also elevated in cells expressing *relish*- targeted sgRNAs, indicating that these togaviruses are also restricted by Relish-dependent antiviral responses (**Figure 6A**). Interestingly, cells expressing either *SHOC2* sgRNA-A or sgRNA-B were significantly more susceptible to infection with ONNV-GFP, SINV-GFP, and VSV- GFP infection (**Figure 6A**). While only cells expressing *SHOC2* sgRNA-A displayed statistically- significant differences in RRV-GFP infection, *SHOC2* sgRNA-B-expressing cells trended towards an increased susceptibility to this virus with an ∼9-fold higher mean in GFP signal than empty vector controls (**Figure 6A**). These results indicate that SHOC2 is a broadly-acting restriction factor for multiple arboviruses in LD652 cells. However, the specific role of SHOC2 in arbovirus restriction requires further investigation.

**Figure 6.**
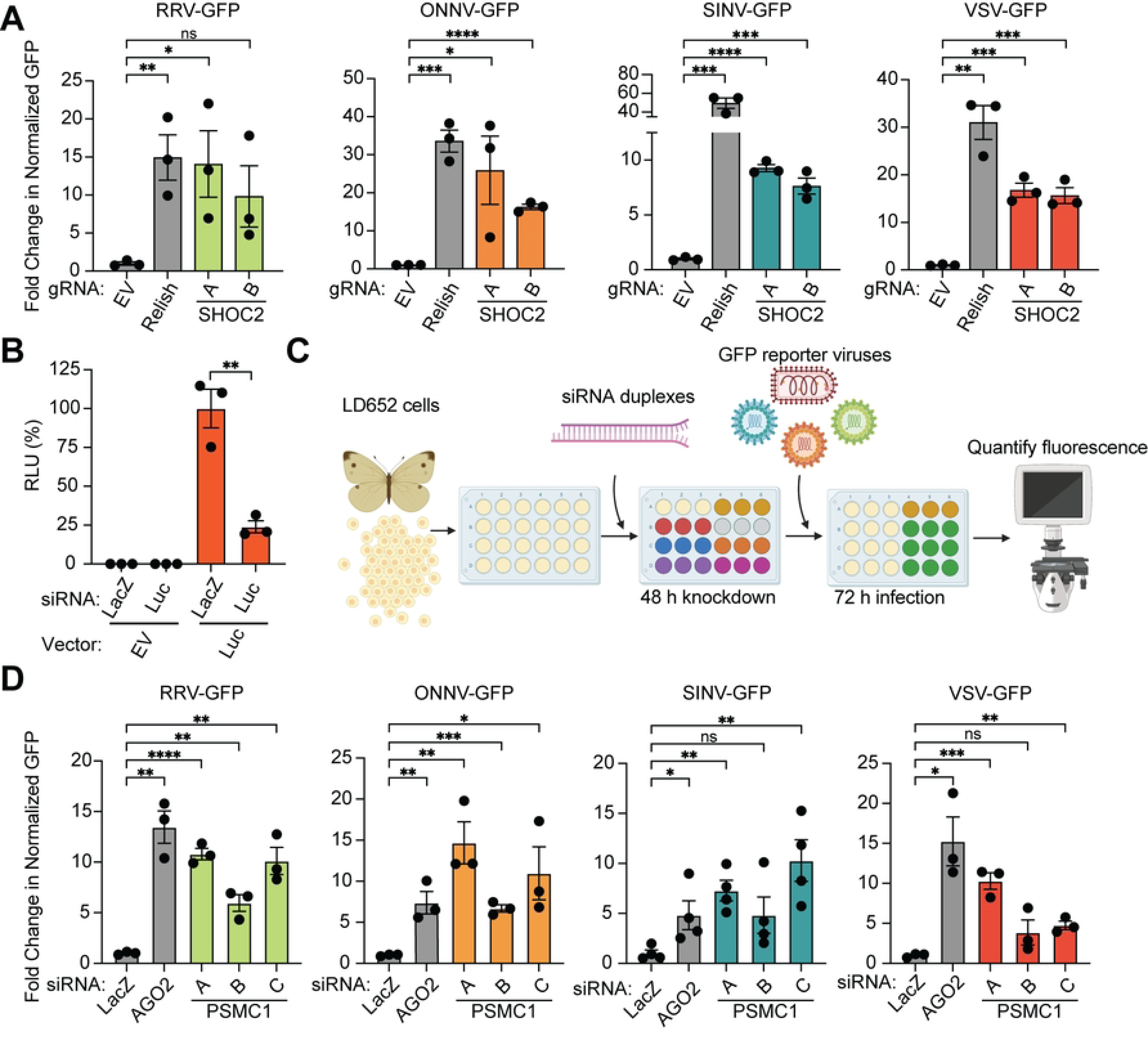
Depletion of IpaH4 substrates SHOC2 and PSMC1 enhances arbovirus replication in LD652 cells. A. Fold-change in normalized viral GFP signals in cells expressing gRNA targeting Relish or SHOC2 relative to empty vector controls 72 hpi. Cells were stained with CellTracker™ dye 72 hpi and imaged to calculate fold-change in normalized GFP signal over empty vector (control) treatments. Data are means ± SD; n=3. Statistical significance was determined with unpaired student’s t-test. **B.** Relative Light Units (RLU) of LD652 lysates from cells transfected with empty vector (EV) or luciferase (Luc)-expressing vectors for 48 and then transfected with siRNA targeting LacZ (negative control) or Luc sequences for 48 h. Data are means ± SD; n=3. Statistical significance was determined with unpaired student’s t-test. **C.** Schematic detailing siRNA knockdown protocol to assess impact of PSMC1 on restricted arbovirus replication in LD652 cells. Image was created with Biorender.com. **D.** Fold-change in normalized viral GFP signals relative to LacZ siRNA (control) treatments. Cells were stained with CellTracker™ dye 72 hpi and imaged to calculate fold-change in normalized GFP signal over LacZ (control) siRNA treatments. Data are means ± SD; n=3. Statistical significance was determined with unpaired student’s t-test; ns=P>0.1234, *=P<0.0332, **=P<0.0021, ***=P<0.0002, ****=P<0.0001.

PSMC1 has been reported to be an essential component of the 19S cap of the 26S proteasome [59]. Consistent with this, our attempts to knock out PSMC1 in LD652 cells with CRISPR-Cas9 techniques resulted in complete cell death after 1-2 rounds of puromycin selection. However, in mammalian systems, transient PSMC1 depletion has been achieved by siRNA knockdown [60]. Therefore, we sought to deplete PSMC1 in LD652 cells in an analogous manner. However, the application of siRNA-based RNAi in *L. dispar* and other lepidopteran cell types has not been well-established. Thus, we took advantage of a prior study that developed guidelines for designing siRNAs to achieve efficient knockdown in another moth species, *Bombyx mori*, as a basis for our siRNA design for use in LD652 cells [61]. To evaluate the efficiency of siRNA- mediated RNAi in LD652 cells, we designed siRNA targeting the coding sequences of *E. coli LacZ* (negative control) and firefly luciferase. Cells were transfected with either an empty vector or a luciferase-encoding pDGOpIE2 plasmid for 48 h and then subsequently transfected with siRNAs targeting transcripts encoding LacZ or luciferase. We then evaluated the relative expression of luciferase using luminescence assays 72 h later. Compared to luciferase signals observed in cells transfected with control LacZ siRNA, there was a significant ∼75% reduction in luminescence signals in cells transfected with siRNA targeting transcripts encoding luciferase (**Figure 6B**), suggesting our siRNA design and transfection strategy was relatively efficient at reducing target gene expression.

We next designed three independent siRNAs targeting *L. dispar PSMC1* sequence in LD652 cells [14] and assessed their relative impact on arboviral replication compared to treatments where control siRNAs targeting LacZ were transfected (**Figure 6C**). As a positive control for arbovirus rescue, siRNAs targeting transcripts encoding argonaute-2 (AGO2), which we have shown to restrict VSV and SINV replication in LD652 cells [62], were also transfected into cells. Compared to LacZ (negative control) siRNA treatments, at least 2/3 *PSMC1*-targeting siRNA transfections resulted in significant increases in viral replication for all four arboviruses (**Figure 6D**). These data indicate that, like SHOC2, PSMC1 may also play a role in restricting arbovirus replication in LD652 cells. Given that PSMC1 is a proteasome subunit, we asked if treatment of LD652 cells with the proteasome inhibitor bortezomib (Bort) would alter their susceptibility to arbovirus infection. Interestingly, addition of Bort to cell culture media 2 hpi resulted in significantly greater viral replication by 72 hpi (**Figure S3AB**). These data suggest that depletion of a proteasome subunit or inhibition of proteasome activity sensitizes LD652 cells to arbovirus infection. However, the mechanism(s) by which PSMC1 and proteasome activity restrict arbovirus replication will require additional studies in the future.

### IpaH4 activity or SHOC2/PSMC1 depletion breaks the restriction of oncolytic virus replication in refractory human cancer cells

After using IpaH4 as a molecular tool to uncover roles for SHOC2 and PSMC1 in viral restriction in insect hosts, we next wanted to examine if these host factors have conserved antiviral roles in mammalian hosts. However, the wild-type arboviruses used in our study are already well-adapted for robust replication in mammalian host cells. Therefore, we sought an alternative approach to examine whether SHOC2 and PSMC1 contribute to virus restriction in mammals.

Oncolytic virotherapy involves the use of viruses to replicate in, and destroy, cancerous cells. These viruses can also invoke anti-tumoral adaptive responses *in vivo*. For example, VSV strains encoding a deletion or arginine substitution of methionine 51 in the VSV matrix (M) protein (VSVΔM51/M51R) are being intensively pursued as a potential oncolytic agent [63–68]. These strains display a relatively safe profile because their mutant M proteins are unable to block cellular gene expression and thus are highly susceptible to innate immune responses (e.g. IFN signaling) that are present in normal cells, but typically defective in transformed cells [63–68]. However, a wide array of human cancer cell lines and tumor types have been shown to be refractory to VSVΔM51/M51R replication, presumably due restriction pathways that are still active in transformed cells [69–73]. This barrier to oncolytic VSV strain replication in refractory cancer types poses a significant challenge to the broad use of these strains for treating diverse malignancies [70, 74]. Given that IpaH4 expression could break wild-type VSV restriction in LD652 cells, we asked whether this bacterial effector could also break restriction of a VSV-M51R strain encoding GFP (VSV-M51R-GFP) in refractory 786-0 human adenocarcinoma cells [71, 75]. Interestingly, transfection of Flag-IpaH4 expression constructs into 786-0 cells led to a significant increase in VSV-M51R-GFP replication compared to empty vector (control) treatments. In contrast, Flag- IpaH4^C339S^ expression did not rescue VSV-M51R-GFP replication, indicating that IpaH4 E3 ubiquitin ligase activity was required to enhance the susceptibility of 786-0 cells to viral replication (**Figure 7AB**). These results also demonstrate that, as in moth cells, IpaH4 can sensitize human cells to VSV infection and demonstrate that bacterial effectors may be useful tools for breaking oncolytic virus restrictions in refractory human cancer cells.

Finally, we asked whether the targets of IpaH4 we identified (SHOC2 and PSMC1) may contribute to virus restriction in human 786-0 cells as observed in moth LD652 cells. To assess the impact of human SHOC2 and PSMC1 on VSV-M51R-GFP restriction, we used three independent siRNAs to deplete 786-0 cells or either SHOC2 or PSMC1 and then assessed VSV- M51R-GFP replication relative to non-targeting (scrambled) siRNA treatments. As a positive control for enhanced VSV-M51R-GFP replication, we also included siRNA treatments targeting *relA*, which encodes a human NF-κB subunit, as NF-κB signaling has been shown to contribute to oncolytic VSV strain restriction in these cells [71]. Compared to control RNAi treatments, knockdown of RelA significantly increased VSV-M51R infection as expected. Interestingly, at least 2/3 and 3/3 siRNAs targeting SHOC2 and PSMC1, respectively, sensitized 786-0 cells to VSV- M51R-GFP infection (**Figure 7CD**). Effective and specific knockdown was confirmed with immunoblotting 786-0 whole cell extracts. Notably, *SHOC2* siRNA-A was not as efficient at knocking down human SHOC2 levels and may explain why we did not detect significant differences in VSV-M51R-GFP replication in those treatments (**Figure 7E**). Collectively, these data suggest that SHOC2 and PSMC1 contribute to virus restriction in human cells and suggest that they may be at least partly responsible for the refractory nature of 786-0 cells to VSV-M51R- GFP replication.

**Figure 7.**
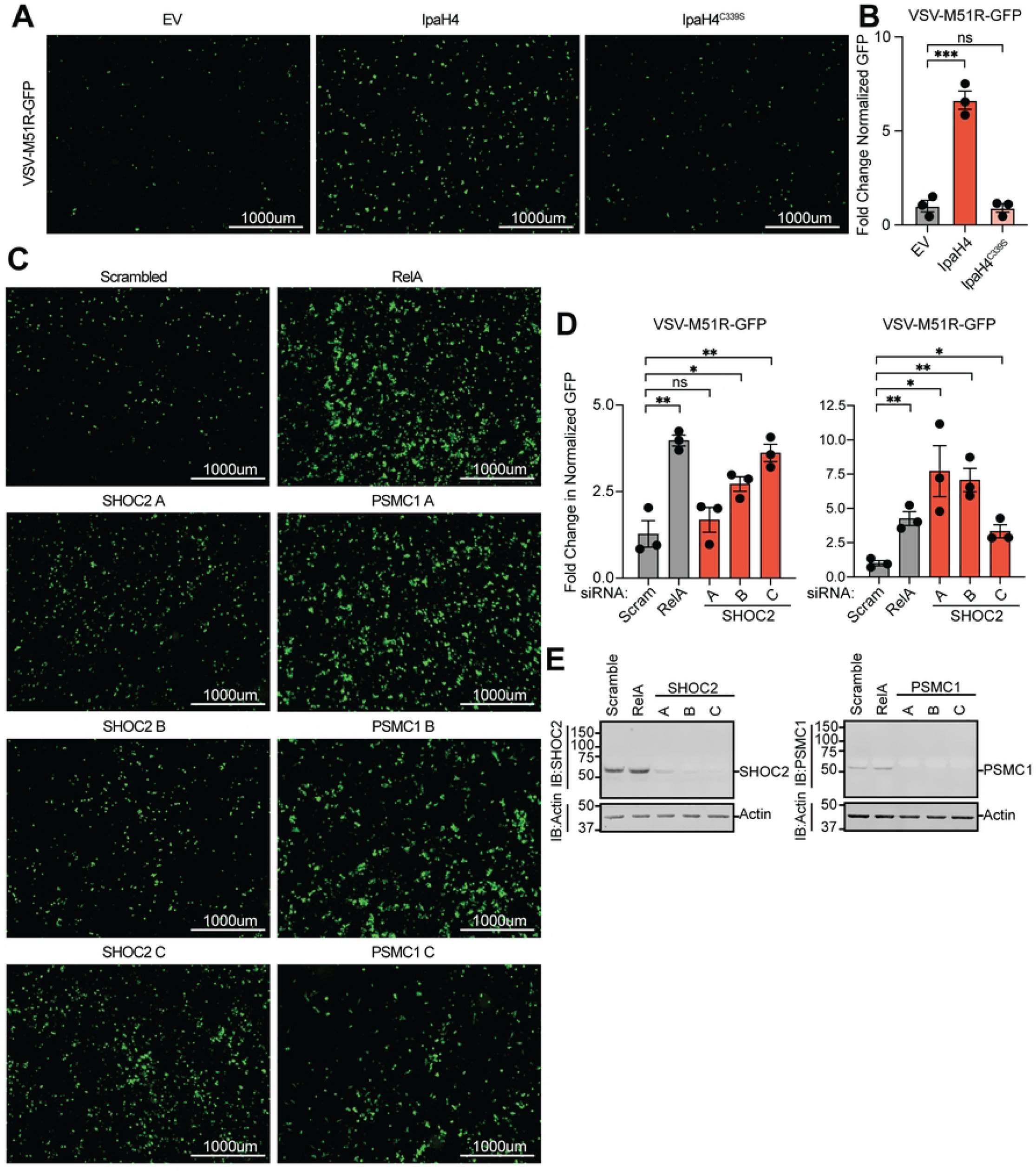
Bacterial effector expression or depletion of effector targets enhances oncolytic virus replication in human cancer cells. A. Representative fluorescence microscopy images (GFP channel) of human 786-0 renal adenocarcinoma cell line infected with VSV-M51R-GFP at 16 hpi. **B.** Fold-change in normalized viral GFP signal relative to empty vector (EV) control from experiments as in **A**. Cells were stained with CellTracker™ Orange dye 16 hpi and imaged to calculate fold-change in normalized GFP signal over EV treatments. Data are means ± SD; n=3. Statistical significance was determined with unpaired student’s t-test. **C.** Representative fluorescence microscopy images (GFP channel) of 786-0 cells infected with VSV-M51R-GFP at 16 hpi after transfection with indicated siRNAs. **D.** Fold-change in normalized viral GFP signal relative to scrambled (control) treatments as in **C**. Cells were stained with CellTracker™ dye 16 hpi and imaged to calculate fold-change in normalized GFP signal over control treatments. **E.** Representative immunoblot of 786-0 whole cell lysate 72 h post-transfection with indicated siRNAs. For **B** and **D**: ns=P>0.1234, *=P<0.0332, **=P<0.0021, ***=P<0.0002, ****=P<0.0001.

## Discussion

Our study illustrates the utility of using abortive virus infections as a screening tool to identify IEPs and the host immunity factors they target. From this work, we identify a variety of bacterially-encoded proteins capable of rescuing individual arbovirus infections, as well as six that are capable of all four viruses screened. Additional analysis into the roles of these effectors showed that three of the effectors with known functions were dependent on their catalytic domains for their viral rescue phenotype. One effector of previously unknown function, Ceg10, was able to be briefly characterized through domain prediction, structure determination, and mutational analysis. Lastly, we established IpaH4 as a strong rescuer for all the arboviruses in our study, and uncovered two evolutionarily conserved targets for this E3 ubiquitin ligase: SHOC2 and PSMC1. These IpaH4 substrates were not only degraded in mammalian culture, but also in invertebrate cells upon expression of wild-type, but not catalytically-inactive, IpaH4. Both SHOC2 and PSMC1 proved important for restricting arbovirus replication in LD652 insect cells and in human 786-0 cells. We envision that our screening methodology can further identify unique IEPs targeting conserved pathways for a variety of pathogen-encoded proteins.

An exciting aspect of our work is that arbovirus replication in moth LD652 cells could be rescued by 30 different bacterial effector proteins, including six (SopB, IpgD, HopAM1, HopT1-2, Ceg10 and IpaH4) that could broadly rescue all four arboviruses examined. Initial characterization of two of these effectors revealed new facets of lipid signaling involved in viral restriction. For example, our studies suggest that acute changes in the phosphorylation status of phosphatidyl inositol (PI) lipids may play a key role in viral restriction in LD652 cells. Not only do both *S. enterica*-encoded SopB and the homologous *S. flexneri*-encoded IpgD rescue arbovirus infection, but inactivation of the phosphatidylinositol phosphatase activities of these bacterial effectors abolished the viral rescue phenotype (**Figure 3C-F**). There have been several reported consequences for the lipid phosphatase activity of SopB and IpgD during bacterial infection including inducing actin remodeling to promote membrane ruffling at the cell surface and bacterial entry [26, 76, 77] and activation of PI3K/AKT signaling to stunt programmed cell death pathways [78, 79]. Interestingly, actin remodeling pathways have been shown to be critical for late stages of togavirus replication wherein these cytoskeletal arrangements are thought to mediate viral envelope protein transport to the cell surface [80]. Furthermore, many togaviruses activate PI3K/AKT signaling during infection and pharmacological inhibition of this signaling pathway has been shown to inhibit togavirus replication in vertebrate cells [81]. Although future mechanistic studies will be required to determine how exactly SopB and IpgD promote arbovirus replication in LD652 cells, it is possible that their phosphatase activity may complement a defect in the ability of these viruses to manipulate actin dynamics and/or regulate PI3K/AKT signaling pathways in these unnatural host cells.

Two unexpected hits from our screen, *P. syringae* effectors HopAM1 and HopT1-2, suggests the exciting possibility that innate immune responses may be conserved between plants and invertebrate animals. *P. syringae* is a plant pathogen that has been an important model for understanding plant immunity to bacterial infection. Recently, HopAM1 was shown to synthesize a novel cADPR variant, v2-cADPR, which has yet to be explored outside of prokaryotic and plant biology [33, 34]. Our data shows that the E191 residue responsible for the ability of HopAM1 to catalyze the v2-cADPR reaction was necessary for arbovirus rescue in LD652 cells (**Figure 3C- F**). This suggests that the v2-ADPR molecule produced by HopAM1 is countering an innate immunity pathway important for antimicrobial responses conserved across phyla. Interestingly, although HopT1-2 lacks clear homology to known enzymes, it has also been previously implicated in anti-bacterial immunity in plants. HopT1-2 can suppress plant nonhost resistance by inhibiting expression of nonhost 1 proteins required for activation of this immune response pathway, although how HopT1-2 achieves this function is unclear [36, 37]. Nonhost resistance responses are driven by nucleotide-binding leucine-rich repeat (NLR) proteins that recognize non-adapted pathogens [36, 37]. NLR proteins are conserved in animals and are also involved in innate immune responses to pathogens [82]. This raises the intriguing possibility that HopT1-2 may target a conserved component of NLR-mediated immunity pathways shared between plant and animal hosts.

By far the largest proportion of effector proteins used to screen arbovirus restriction are encoded by *Legionella pneumophila*, a bacteria species that utilizes a Type IV secretion system to deliver ∼350 effector proteins into the host cells. While many of these effectors exhibit unique enzymatic activities and play different roles during infection, most remain uncharacterized [83]. We found the uncharacterized *L. pneumophila*-encoded Ceg10 protein to reproducibly break viral restriction in LD652 cells. Interestingly, the conserved catalytic cysteine, Cys-159 (C159), is nitrosylated in two of our three structural data sets, indicating that this cysteine is highly reactive and particularly susceptible to oxidation by agents such as nitric oxide. Recent studies indicate that S-nitrosylation of the *Vibrio cholera* virulence regulator AphB suppresses the enzymatic activity and alters virulence gene expression [84]. To date, of the >215,000 structures available on the Protein Data Bank, only 43 deposits display an S-nitrosothiol group, highlighting the unique nature of this modification. Furthermore, to our knowledge, Ceg10 is the first structurally characterized S-nitrosylated *Legionella* effector. We confirmed the importance of this residue by mutating C159 to serine and demonstrated that mutant Ceg10 is not capable of breaking viral restriction in L652 cells. These data suggest that Ceg10 targets a highly conserved immunity evasion factor that may be important for controlling both viral and bacterial infections. Furthermore, it reveals a potential mechanism for host defense against Ceg10 via S-nitrosylation and subsequent inactivation.

While determining how SopB, IpgD, HopAM1, HopT1-2 and Ceg10 rescue virus replication in moth cells will be an exciting avenue of future research, we sought to identify novel points of viral restriction by exploring the activity of IpaH4 in greater detail given our prior interest in IpaH family members [46–48]. Through independent UBAIT- and Y2H-based screening methods, we identified host SHOC2 and PSMC1 proteins as conserved IpaH4 substrates in moth and human cells. SHOC2 is a leucine-rich repeat scaffold protein that forms a ternary complex with MRAS and PP1C that activates rapidly accelerated fibrosarcoma (RAF) kinases [85, 86]. Although the specific role of SHOC2 during virus infection is unclear, it has recently been shown to activate ERK/STAT signaling in response to bacterial flagellin in shrimp hosts [87], implying that SHOC2 may indeed have conserved roles in combating bacterial and viral pathogens. Although it is unknown if SHOC2 restricts *S. flexneri* infection, this bacterium lacks flagella. Moreover, recognition of flagella cannot explain how SHOC2 contributes to arbovirus restriction, suggesting that different pathogens can activate SHOC2-mediated immunity through distinct mechanisms. These findings highlight the strength of our system for identifying pathogen- encoded modulators of key innate immunity pathways that are likely relevant to both bacterial and viral pathogens.

PSMC1 is an essential component of the 19S cap of the proteasome functioning as one of six ATPase subunits [88]. In yeast, PSMC1 (RPT2) controls the gate of the proteasome and regulates the opening through an ATP-binding motif, allowing for substrates to enter [59]. Mutations within the ATP-binding motif in yeast PSMC1 was lethal and purified proteasomes exhibited substantially less peptidase activity [89]. Data suggests a similar role of PSMC1 in human proteasomes, although it remains unclear if other ATPase subunits are involved in gate-regulating activity [88, 90]. We show that both knockdown of PSMC1 and suppression of proteasome activity alleviate the restricted arbovirus replication seen in LD652 cells. It is attractive to speculate that the proteasome degrades proteins required for the viral lifecycle in these cells, and that IpaH4 suppresses this activity resulting in viral replication. However, it is entirely possible that PSMC1-mediated virus restriction may be proteasome-dependent or -independent given the growing evidence that proteasome subunits play roles outside the proteasome complex [57, 91, 92]. Future research will be needed to address the sophisticated interaction between viral restriction and proteasome functions in the invertebrate host.

Given that the arboviruses in our study robustly replicate in human cells, we suspected that it may be more difficult to identify roles for human PSMC1 and SHOC2 in the restriction of these viruses. Thus, we sought to recapitulate a similar restricted arbovirus replication system as found in LD652 cells but in a human cell background. To do this, we took advantage of the fact that the oncolytic virotherapy strain, VSV-M51R, poorly replicates in human renal adenocarcinoma 786-0 cells due to an inability of this mutant virus to inhibit innate immune responses [63–68]. We showed that 786-0 cells became sensitize to VSV-M51R infection after either overexpression of IpaH4 or knockdown of IpaH4 substrates, PSMC1 and SHOC2 (**Figure 6**). This raises the interesting possibility that either bacterial effector function or inactivation of the host targets of effectors might enhance the replication of oncolytic viruses in cancer cell types normally refractory to these viruses. Importantly, both SHOC2 and PSMC1 are often up-regulated in human malignancies have been linked to cellular transformation and tumor metastasis [93–95]. Indeed, gain-of-function SHOC2 mutations have been linked to RASopathies, a group of clinical pathologies associated with dysregulated RAS/MAPK signaling such as Noonan syndrome and patients with these syndromes are predisposed to certain cancers [96–98]. Moreover, elevated SHOC2 and PSMC1 expression has been associated with poorer clinical outcomes among cancer patients [95, 98]. Given the common upregulation of SHOC2 and PSMC1 in transformed cells, they may be particularly relevant to the reported refractory nature of some tumors to oncolytic VSV strains replication [70, 74]. Our work opens up the exciting possibility that expression of IpaH4 (or inhibition of the host factors IpaH4 targets) may provide a novel strategy for increasing oncolytic VSV strain replication in refractory tumors that, in turn, may improve virotherapy efficacy.

## Limitations of the study

Our study utilizes a less developed lepidopteran cell system where reagents (e.g. antibodies) that could be used to validate SHOC2 or PSMC1 knockouts/knockdowns in *L. dispar* cells are not currently available. Furthermore, attempts to use antibodies raised against mammalian SHOC2 or PSMC1 failed to show cross reactivity in LD652 whole cell extract. Thus, we either used alternative methods for validation of knockdown procedures in LD652 cells (ex. Figure 5C) and/or multiple, independent sgRNAs or siRNAs to confirm SHOC2- and PSMC1-related phenotypes in insect cells. Furthermore, because our focus here was on identifying host factors that contribute to virus restriction, we have not yet determined if SHOC2 or PSMC1 are targets of IpaH4 during intracellular *S. flexneri* infection. These future studies will be important for understanding the impact of IpaH4-eukaryotic host interactions to bacterial replication and pathogenesis.

## Materials and Methods

### Cell Lines and Cell Culture

Mammalian cell lines were maintained at 37°C in 5% CO_2_ atmosphere. HEK293T and BHK-21 cells were cultured in DMEM supplemented with 10% FBS. 786-0 cells were cultured in RPMI supplemented with 10% FBS. BSC-40 cells were cultured in MEM supplemented with 5% FBS. All media additionally contained 1% non-essential amino acids, 1% L-glutamine, and 1% antibiotic/antimycotic (Gibco). LD652 cells were cultured as described previously in a 1:1 mixture of Grace’s Insect Media (Sigma) and Ex-Cell 420 (Sigma) at 27°C under normal atmospheric conditions [12].

### Viruses

Stock preparation and culture of recombinant VSV and SINV was performed as previously described [12]. VSV-M51R-eGFP was obtained from Dr. Doug Lyles (Wake Forest University). RRV-GFP (strain T48), ONNV-GFP (strain SG650) constructs were obtained from Dr. John Schoggins (UTSW Medical Center). VSV, SINV, RRV, and ONNV stocks were amplified using low MOI conditions in BHK-21 cells. Viruses were collected from supernatants using ultracentrifugation (22,000 rpm, 2 h, 4°C) and titrated on BSC-40 using fluorescent plaque/foci assays as described [12].

Viral infections were incubated for 2 h in serum free media (DMEM for mammalian cells or Sf-900 II for invertebrate cells; **Table S4**) before the inoculum was replaced with complete media for the remainder of the infection. Where indicated, complete media containing 0.05 µg/mL ActD or 50 nM Bort was added for the remainder of the infection.

### Plasmid Constructs for Mammalian and Insect Cell Expression

The bacterial gene library was assembled from a previously generated pENTR library of genes [99]. Additional bacterial effector genes were added to the pENTR library by Gateway® Cloning using BP Clonase II (Invitrogen; **Table S4**), following the manufacturer’s instructions. All pENTR effector gene were verified by Sanger sequencing. The entire pENTR effector library was then transferred into the pIB/V5-His vector (Invitrogen; **Table S4**) using LR Clonase II (Invitrogen; **Table S4**), following manufacturer’s instructions.

N-terminal Flag-tagged versions of IpgD codon-optimized for expression in insect cells [IpgD(CO)], SopB, Ceg10, HopT1-2, and HopAM1 and C-terminal Flag-tagged IpaH4 were generated by PCR amplification using primers containing Flag sequences and flanking SacII / PacI off pIB/V5-His templates using iProof DNA polymerase (Bio-Rad; **Table S4**) and cloned into the pDGOpIE2 vector. The pDGOpIE2 vector was synthesized by Gene Universal and contains an OpIE2 promoter, multiple cloning site, polyA sequence and puromycin resistance cassette. A plasmid map of the pDGOpIE2 vector can be found in **Figure S1**. Catalytically-inactive mutants of IpaH4, SopB, and IpgD were constructed using site-directed mutagenesis by PCR amplification using Q5 DNA polymerase (NEB; **Table S4**).

N-Terminal GFP-IpaH4 was constructed through gateway cloning into pEGFP-C2 (for mammalian expression), then PCR amplified using primers introducing SacII / PacI sites and cloned into pDGOpIE2 (for insect cell expression).

N-terminal Flag human PSMC1 and SHOC2 were generated via Gateway cloning into pCDNA3, (for mammalian cell expression), then PCR amplified using primers introducing SacII / PacI sites and cloned into pDGOpIE2 (insect cell expression).

N-terminal Flag *L. dispar* PSMC1 and SHOC2 constructs were generated by RT-PCR amplification from total RNA isolated from LD652 cells using Superscript III reverse transcriptase (Thermo Fisher), iProof DNA polymerase, and primers containing SacII/PacI sites. SacII/PacI constructs were cloned into a modified pCDNA3 vector containing SacII/PacI sites [12] or pDGOpIE2.

### Immunoblotting

Protein extracts were diluted in 5X SDS-PAGE loading buffer then boiled at 95°C for 10 min. Samples were subjected to SDS-PAGE electrophoresis at 125 V for approximately 1.5 h. Separated proteins were transferred to nitrocellulose membranes in either 1X transfer buffer (BioRad; **Table S4**) at 1300 mA at 25°C, or in Towbin Buffer (BioRad; **Table S4**) at 100 V at 4°C for 100 min. Membranes were blocked in Odyssey Blocking Buffer (LI-COR; **Table S4**) for 1 h at 25°C. Membranes were blotted with primary antibody overnight at 4°C, with actin serving as a loading control unless otherwise stated. After three, 5 min washes with PBS-T (PBS, 0.1% Tween; **Table S4**), membranes were incubated in secondary antibody conjugated to an IRDye (LI-COR; **Table S4**) for 1 h followed by two, 5 min washes in PBS-T and one 5 min PBS wash. Membranes were then imaged with an Odyssey Fc Imager (LI-COR; **Table S4**).

### Fluorescence Microscopy

Cells were stained with 200 µL of serum free media containing CellTracker™ Orange (Invitrogen; **Table S4**) dye at specified concentrations for 30 min followed by replacement with 1 mL PBS. Cells were imaged using a 2X objective on an EVOS-FL fluorescence microscope (Thermo Fisher; **Table S4**) using RFP and GFP cubes. Each condition had 3 replicate wells, and two images/well were collected for analysis. Image analysis was conducted using Fiji (NIH) to quantify the percent area of each field of view containing GFP and RFP signal. Positive signal was determined by using uninfected wells lacking CellTracker™ stain to set a minimum threshold in both GFP and RFP channels that was applied to the remaining dataset. Only signal above the set threshold was highlighted and percent area of each field of view with fluorescent signal was measured using the Fiji analyze particles function. GFP signals were then divided by the percent area of RFP to give normalized GFP signal. Fold change in GFP signals were calculated by dividing normalized GFP values of experimental treatments with normalized GFP values in control treatments (indicated in each figure).

### Structural determination of Ceg10

To determine the structured core of Ceg10 (NCBI accession number YP_094338.1), primers were designed to clone Ceg10 into pET28b using restriction enzymes NdeI and NotI by HiFi DNA Assemby kit (NEB; **Table S4**), producing the His_6x_-TEV-Ceg10 vector. *E. coli* BL21 cells transformed with His_6x_-TEV-Ceg10 were grown in LB medium supplemented with kanamycin (50 mg/ml). After cultures reached an OD_600_ of 0.4-0.5, protein expression was induced with 0.2 mM IPTG overnight at 18°C. Cells were lysed using an Emulsiflex C5 (Avestin) in buffer containing 50 mM Tris-base, 500 mM sodium chloride (NaCl), 10 mM imidazole, 1 mM phenylmethylsulfonyl fluoride (PMSF), 0.5 mM tris(2-carboxyethyl)phosphine (TCEP), and SIGMAFAST™ Protease Inhibitor Cocktail Tablets at pH 7.0. The fusion protein was affinity purified using Ni-NTA agarose (Qiagen; **Table S4**), eluted with 50 mM Tris-base, 150 mM NaCl, 250 mM imidazole, and 0.1 mM PMSF, and dialyzed overnight into 50 mM Tris-base, 150 mM NaCl, and 1 mM magnesium chloride (MgCl_2_) at 4°C. Limited proteolysis of Ceg10 was carried by digesting 6.4 μM His_6x_-TEV-Ceg10 with 0.89 μM Trypsin (Sigma) in PBS. After 30 min at 25°C, the reaction was quenched by the addition of 1 mM PMSF and the molecular weight of the core was determined by intact mass spectroscopic analysis (UTSW Proteomic Core), revealing a highly abundant peptide (T55-R287) corresponding to residues Thr-55-Arg287 (Ceg10^TR^) in Ceg10. Primers were designed to clone Ceg10^TR^ pET28b as described above to produce the His_6x_-TEV-Ceg10^TR^ vector. *E. coli* BL21 cells transformed with His_6x_-TEV-Ceg10 were grown in LB medium supplemented with kanamycin (50 μg/ml). After cultures reached an OD600 of 0.4-0.5, protein expression was induced with 0.2 mM IPTG overnight at 18°C. Cells were lysed using an Emulsiflex C5 (Avestin) in buffer containing 50 mM Tris-base, 500 mM NaCl, 10 mM imidazole, 1 mM PMSF, 0.5 mM TCEP, and SIGMAFAST™ Protease Inhibitor Cocktail Tablets at pH 7.4. The fusion protein was affinity purified using Ni-NTA agarose (Qiagen), eluted with 50 mM Tris-base, 150 mM NaCl, 250 mM imidazole, 0.5 mM TCEP and 0.1 mM PMSF, and dialyzed overnight into 50 mM Tris-base, 150 mM NaCl, and 0.5 mM TCEP at 4°C. The 6XHis-tag was removed in the presences of 1 mM EDTA by digestion with His- tagged Tobacco Etch Virus (TEV) protease (1:50 TEV:protein ratio). After 16 h at 25°C, MgCl_2_ (5 mM final) was added and TEV protease was removed by passing the cleave reaction over 3 ml of Ni-NTA agarose, followed by washing with 20 ml of 10 mM Tris-base, 50 mM NaCl, and 1 mM TCEP at pH 8.0. Ceg10^TR^ was concentrated to 2.5 ml using a 10,000 MWCO Amicon 50 spin concentrator, subjected to centrifugation at 40,000 rpm (TLA55 rotor), and further purified using a Superdex highload 200 16/600 gel filtration chromatography column (Cytiva Life Sciences). Fractions containing Ceg10^TR^ were concentrated of 20 mg/ml and filtered with a 0.22 μm Durapore® membrane filter.

Native crystals were grown by the sitting drop vapor diffusion method at 4°C in 96-well Intelliplate trays using a 1:1 ratio of protein/reservoir solution containing 1.5 M sodium phosphate monobasic, 0.5 M potassium phosphate dibasic, 10 mM sodium phosphate dibasic/citrate buffer at pH 4.2 and were cryoprotected with 25% ethylene glycol. Native crystals diffracted to a minimum Bragg spacing (dmin) of 1.70 Å and exhibited the symmetry of space group C2221 with cell dimensions of a = 86.0 Å, b = 112.3 Å, c = 55.2 Å, and contained one Ceg10^55-287^ per asymmetric unit. Crystals of S-nitrosylated Ceg10^55-287^ were grown by the sitting drop vapor diffusion method at 4°C in 96-well Intelliplate trays using a 1:1 ratio of protein/reservoir solution containing 20% PEG 1,000, 0.1 M Tris pH 7.0 and were cryoprotected with 25% ethylene glycol. S-nitrosylated Ceg10^55-287^ crystals diffracted to a minimum Bragg spacing (dmin) of 1.40 Å and exhibited the symmetry of space group P2_1_2_1_2 with cell dimensions of a = 103.4 Å, b = 115.8 Å, c = 40.9 Å, and contained two Ceg10^55-287^ per asymmetric unit.

Crystals of Ta_6_Br_12_^2-^ derivatized S-nitrosylated Ceg10^55-287^ were grown by the sitting drop vapor diffusion method at 4°C in 96-well Intelliplate trays using a 1:1 ratio of protein/reservoir solution containing 10% PEG 8,000, 10% PEG 1,000, 0.2 M MgCl_2_, 0.1 M sodium acetate pH 5.0 and were cryoprotected with 25% ethylene glycol. Derivatized S-nitrosylated Ceg10^55-287^ crystals diffracted to a minimum Bragg spacing (dmin) of 1.52 Å and exhibited the symmetry of space group P2_1_2_1_2 with cell dimensions of a = 103.6 Å, b = 116.4 Å, c = 40.8 Å, and contained two Ceg10^55-287^ as well as two Ta_6_Br_12_^2-^ clusters per asymmetric unit. All diffraction data were collected at beamline 19-ID (SBC-CAT) at the Advanced Photon Source (Argonne National Laboratory, Argonne, Illinois, USA) and processed in the program HKL-3000 [100] with applied corrections for effects resulting from absorption in a crystal and for radiation damage [101, 102], the calculation of an optimal error model, and corrections to compensate the phasing signal for a radiation-induced increase of non-isomorphism within the crystal [103, 104].

Phases were obtained from a single wavelength anomalous dispersion (SAD) experiment using the Ta_6_Br_12_^2-^ derivatized S-nitrosylated Ceg10^55-287^ with data to 1.52 Å collected at the LIII-edge of tantalum. Twelve tantalum sites were located by the program AutoSol, part of the Phenix package [105], and an initial model containing 78.5% of all Ceg10^55-287^ residues was automatically generated. This model was used for molecular replacement phasing of the native Ceg10^55-287^ and the S-nitrosylated Ceg10^55-287^. Completion of these models was performed by multiple cycles of manual rebuilding in the program Coot [106]. Positional and isotropic atomic displacement parameter (ADP) as well as TLS ADP refinement was performed using the program Phenix with a random 5.0% of all data set aside for an R_free_ calculation. Data collection and structure refinement statistics are summarized in Table S2.

For determining the electrostatic potential of Ceg10 and Ceg10^TR^ the APBS plugin for PyMOL (PyMOL Molecular Graphics System, Version 2.0 Schrödinger, LLC) was used.

### Luciferase Assay

Luciferase assays with were performed as described with minor modifications [12, 22]. Briefly, at indicated times post infection/transfection, cells were washed in PBS, pelleted by brief centrifugation, and lysed in reporter lysis buffer (Promega; **Table S4**). WCE was spotted in a 96-well dish and luciferase activity was measured in arbitrary light units (LU) using a FLUOstar Omega plate reader (BMG Labtech).

### RNAi and Viral Infection in Cell Culture

Each 21-nucleotide siRNA sequence was designed based on gene prediction using LD652 genome [14] following rules established by others RNAi experiments in *Bombyx mori* [61]. siRNAs were designed against target sites that were at least 75 nts downstream of the initiation codon of each target gene, were 19 nts in length, and had a GC content of 35-55% [61]. Targets were chosen such that the sense siRNA start nucleotide was preceded by “AA” sequence and the antisense siRNA ended on an A or U [61]. Transient siRNA knockdown was achieved by forward transfection of LD652 with Cellfectin II (Gibco; **Table S4**) according to the manufacturer’s protocol for 48 h prior to infection. Cells were then infected with virus for 72 h prior to imaging-based quantification of infection.

### Y2H Screens

Y2H screening was conducted by Hybrigenics Services (Boston, MA). The coding sequence for IpaH4 was PCR amplified from Flag-IpaH4 pCDNA3 vectors and cloned into pB66 as a C-terminal fusion to the Gal4 DNA-binding domain (but without the Flag tag) creating Gal4-IpaH4 pB66. The construct was used as a bait for two independent screens with a random-primed human lung cancer cDNA library constructed into pP6. Clones were screened at 4-fold complexity of the library using mating approach with YHGX13 (Y187 ade2-101::loxP-kanMX-loxP, mata) and CG1945 (mata) yeast strains. His+ colonies were selected on a medium lacking tryptophan, leucine and histidine. Prey fragments of the positive clones were amplified by PCR and sequenced at their 5’and 3’ junctions. The resulting sequences were used to identify the corresponding interacting proteins in the GenBank database (NCBI).

### UBAIT

For UBAIT assay to capture unknown substrates from cell lysates, LD652 cells were grown to confluency in three 15 cm dishes. Cells were washed with cold PBS and lysed in 300 µL 1x Ubiquitination buffer (Boston Biochem; **Table S4**) containing 1% Triton X-100, 1x Protease inhibitor (Sigma), and 1mM DTT. The lysate was centrifuged for 10 min at 13,000 g and 4°C. The supernatant was transferred to a new tube before adding 25 µg of the GST-IpaH4-3xFlag-ub construct and incubating for 1 h at 4°C with rotation. His-UbE1 (100 nM), His-UbcH5b (2000 nM), ATP (1 mM), MgCl (1 mM) and 1x Ubiquitination buffer (Boston Biochem; **Table S4**) were added and the reaction was incubated at 30°C for 10 min. Following incubation, 300 µL TBS buffer (25 mM Tris-HCl pH 7.4, 150 mM NaCl, 1 mM DTT) and 30 µL of washed GST beads were added. The reaction was incubated at 4°C for 2 h with rotation. The reaction was transferred to a column and the beads were washed four times in TBS containing 0.5% Triton-X 100 and two times in TBS before eluting in 150 µL reduced glutathione pH 8.0. Beads were centrifuged and the supernatant was transferred to a new tube followed by addition of 0.25% SDS (final concentration) and 5 mM DTT (final concentration). The solution was boiled for 5 min at 95 °C before adding 1.2 mL TBS and 20 µL M2-Flag beads (Sigma; **Table S4**) and incubated at 4°C for 2 h with rotation. Flag beads were then washed four times in TBS containing 0.5% Triton X-100 and twice with TBS. Finally, 35 µL of 95°C SDS-PAGE loading buffer was added, and the beads were boiled an additional 5 min at 95°C. Samples were subjected to SDS-PAGE and Coomassie stained. To identify captured proteins, a lane of the gel above the unmodified IpaH4-UBAIT band was excised and proteins were digested in-gel with trypsin and run on a Q Exactive MS platform at the University of Texas Southwestern Medical Center Proteomics Core.

### Degradation Assay

100,000 HEK-293T cells were co-transfected with 150 ng GFP/GFP-IpaH4/GFP-IpaH4^C339S^ and 350 ng target vectors for 48 h using FuGENE (Promega; **Table S4**). Cells were harvested in RIPA buffer containing 100 µM PMSF and protease inhibitor cocktail (Abcam; **Table S4**). Protein extracts were then subjected to SDS-PAGE and subsequent immunoblotting as indicated in each figure.

### Protein Purification

To obtain C-terminal 5XHis-tagged PSMC1, human *PSMC1* was cloned into pET21a by restriction cloning with NdeI / XhoI and expressed in *E. coli* BL21in the presence of ampicillin. After the cultures reached OD_600_ 0.7, protein expression was induced with 0.5 mM IPTG at 18 °C overnight. Cells were lysed using an Emulsiflex C5 (Avestin) in buffer containing 25 mM HEPES, 100 mM NaCl, 100 mM KCl, 10% glycerol, 10 mM MgCl_2_, 0.5 mM EDTA, 1 mM DTT, and 10 mM Imidazole (HBSi; **Table S4**) with the final pH at 7.5. The fusion protein was affinity-purified using TALON Metal Affinity Resin (Takara Biosciences; **Table S4**) and eluted in buffer containing 25 mM HEPES, 100 mM NaCl, 100 mM KCl, 10% glycerol, 10 mM MgCl_2_, 0.5 mM EDTA, 1 mM DTT, and 500 mM Imidazole. The protein was dialyzed using a 10,000 MWCO Slide-A-Lyzer™ G3 dialysis cassette (Thermo; **Table S4**) in 1 L of buffer containing 25 mM HEPES, 100 mM NaCl, 100 mM KCl, 10% glycerol, 10 mM MgCl_2_, 0.5 mM EDTA, 1 mM DTT.

To obtain N-terminal 6XHis-MBP-TEV-Flag-tagged SHOC2, *E. coli* carrying pET28a-SHOC2 were grown in LB media supplemented with ampicillin. After the cultures reached OD_600_ 0.7, protein expression was induced with 0.5 mM IPTG at 17°C overnight. Cells were lysed by sonication with 5 second pulses and 20 second intervals for 50 min at 4°C in buffer containing 50 mM Tris-HCl, 150 mM NaCl, 1 mM TCEP, 1 mM PMSF, and 20 mM Imidazole (HBSi) with the final pH at 8. The lysate was then subjected to centrifugation at 10,000 x g for 30 min at 4°C. The fusion protein was affinity-purified using TALON metal affinity resin (Takara Biosciences) and eluted in buffer containing 50 mM Tris-HCl, 150 mM NaCl, 1 mM TCEP, 0.1 mM PMSF, and 250 mM Imidazole. The protein was concentrated using a 30,000 MWCO Amicon 50 spin concentrator then dialyzed using a 10,000 MWCO Slide-A-Lyzer™ G3 dialysis cassette (Thermo) in buffer containing 25 mM Tris-HCl and 75 mM NaCl. Flag-SHOC2 was cleaved from 6X-His-MBP by incubating 37 mg purified protein with 0.65 mg TEV protease at 4°C overnight followed by affinity purification using Nickel affinity resin (Qiagen) where cleaved Flag-SHOC2 was collected in the flow-through.

Recombinant GST-IpaH4-3XFlag for UBAIT experiments was obtained as previously described for other IpaH proteins [107, 108]. Briefly, GST-IpaH4-3XFlag was cloned into pGEX6P-1 with a 3XFlag peptide followed by the coding sequence for ubiquitin using Gibson Cloning (NEB; **Table S4**). Sequence-verified constructs were transformed into *E. coli* BL21 cells, induced, and recombinant protein was purified using glutathione sepharose beads (GE Healthcare Life Sciences).

Recombinant GST-IpaH4 for *in-vitro* experiments was expressed and purified as previously described for other IpaH proteins [107, 108]. Briefly, GST-IpaH4 was cloned into pGEX6P-1, transformed into *E. coli* BL21 cells, induced, and recombinant protein was purified using glutathione sepharose beads (GE Healthcare Life Sciences; **Table S4**).

### In-Vitro Ubiquitination Assay

*In vitro* ubiquitination reactions were performed in 50 mM HEPES pH 7.5, 150 mM NaCl, 20 mM MgCl_2_, and 10 mM ATP (47). Components were mixed as indicated at the following concentrations 1 µM UbE1 (E1), 5 µM UbcH5b (E2), 500 nM-10 µM GST-IpaH or GST-IpaH4^C339S^, 50 µM ubiquitin, and 5 µM His-tagged PSMC1, or 5 µM Flag-tagged SHOC2. Reactions were incubated for 18 h at 30°C before the addition of 2X loading buffer containing β-mercaptoethanol (BME).

Samples were then boiled for 10 min at 95°C, subjected to SDS-PAGE, and immunoblotted with anti-GST, anti-Ubiquitin, anti-His or anti-Flag antibodies.

### Statistical Analysis

Graphs and charts were presented as mean values ± standard error of mean (SEM) with individual data points for each independent experiment shown. At least three independent experiments were conducted for all quantitative experiments shown. All statistical analyses were performed with Prism software v10.0.2 (GraphPad) and statistical tests used are indicated in respective figure legends. Statistical significance (*P*<0.05) between compared groups is indicated in figures as either: ns (not significant), *=P<0.05, **=*P*<0.01, ***=*P*<0.001, ****=*P*<0.0001.

## Acknowledgements

We would like to thank Drs. John Schoggins and Andrew Sandstrom (UT Southwestern Medical Center) and members of the Gammon and Alto laboratories for helpful discussions. We also thank Dr. Don Jarvis (University of Wyoming) for providing the pIE1-Cas9-SfU6-sgRNA-Puro vector. We thank the UT Southwestern Medical Center Proteomics Core for assistance with aspects of experimental execution. This work was supported by grants to DBG from the NIH (1R35GM137978-01 and 1R21AI169558-01A1) and by funding to DBG from the UTSW Endowed Scholars Program. NMA was supported by NIH NIAID R01AI083359, The Welch Foundation (I- 1704) and The Burroughs Welcome Fund (1011019). This research was also supported with training grant funding from the NIH to AE, NSB, DBH (T32 AI007520).

## Author Contributions

**Aaron Embry:** Conceptualization; Data Curation; Formal Analysis; Funding Acquisition; Investigation; Methodology; Validation; Visualization; Writing – Original Draft; Writing – Review & Editing. **Nina S. Baggett:** Conceptualization; Data Curation; Funding Acquisition; Investigation; Methodology; Validation; Visualization; Writing – Original Draft; Writing – Review & Editing. **David B. Heisler:** Conceptualization; Data Curation; Investigation; Methodology; Resources; Writing – Original Draft; Writing – Review & Editing. **Addison White:** Data Curation; Investigation; Writing – Review & Editing **Maarten F. de Jong:** Data Curation; Investigation; Resources. **Benjamin L. Kocsis:** Investigation; Resources. **Diana R. Tomchick:** Data Curation; Formal Analysis; Investigation; Methodology; Visualization; Writing – Original Draft. **Neal M. Alto**: Conceptualization; Formal Analysis; Supervision; Funding Acquisition; Methodology; Project Administration; Visualization; Writing – Review & Editing. **Don B. Gammon**: Conceptualization; Data Curation; Validation; Formal Analysis; Supervision; Funding Acquisition; Investigation; Methodology; Project Administration; Visualization; Writing – Original Draft; Writing – Review & Editing.

## Data Availability

All relevant data are within the manuscript or supporting data. Ceg10 unmodified and S-nitroylated structures (.pdb files) are included with this manuscript.

## Supplementary Tables

**Supplemental Table 1. Bacterial effector library screening results. 1A.** Fold-change in GFP or luciferase reporter signals for each reporter arbovirus after transfection of indicated expression construct.

**Supplemental Table 2. Data collection and refinement statistics, Ceg10 structures.**

**Supplemental Table 3. Identification of IpaH4 substrates.** Screening putative substrates of IpaH4.

**Supplemental Table 4. Key resources and reagents. Supplementary Figure Legends**

**Figure S3.** Proteasomal activity plays a role in restricting arbovirus replication in LD652 cells. A. Representative fluorescence microscopy images (GFP channel) of LD652 cells 72 hpi with the indicated GFP reporter strains that were treated with DMSO (vehicle) or 50 nM Bortezomib (Bort). DMSO or Bort was added 2 hpi. B. Fold-change in normalized GFP signals in Bort-treated cultures relative to DMSO treatments. Cells were stained 72 hpi with CellTracker™ Orange dye (not shown) and imaged in GFP and RFP channels to calculate fold-change in GFP signal after normalization of cell number using CellTracker™ (RFP) channel signals. Data in B are means ± SD; n=3. Statistical significance was determined with unpaired student’s t-test; ns=P>0.1234, *=P<0.0332, **=P<0.0021, ***=P<0.0002, ****=P<0.0001.

**Figure S1.** Generation of insect expression vector pDGOpIE2. A. Snapgene vector map of pDGOpIE2 vector and features of interest. B. Complete sequence of pDGOpIE2 vector.

**Figure S2.** Identification of putative targets of bacterial E3 ubiquitin ligase IpaH4. A. Table summarizing results of two independent Y2H screens using a human prey library and three independent ubiquitin-activated interaction trap (UBAIT) assays using LD652 cell lysate. Hits were then analyzed via Blastp to determine percent identity to their Homo sapiens ortholog. N.D. = Not determined; either no ortholog found or no significant homology (as determined by BLAST). (B) Representative immunoblot of degradation assays for Flag-tagged human proteins following 48 h co-expression in HEK293T cells with GFP, IpaH4 (WT) or catalytic mutant GST-IpaH4C339S (C339S).

